# Regulation of human development by ubiquitin chain editing of chromatin remodelers

**DOI:** 10.1101/2020.01.23.917450

**Authors:** David B. Beck, Mohammed A. Basar, Anthony J. Asmar, Joyce Thompson, Hirotsugu Oda, Daniela T. Uehara, Ken Saida, Precilla D’Souza, Joann Bodurtha, Weiyi Mu, Kristin W. Barañano, Noriko Miyake, Raymond Wang, Marlies Kempers, Yutaka Nishimura, Satoshi Okada, Tomoki Kosho, Ryan Dale, Apratim Mitra, Ellen Macnamara, Undiagnosed Diseases Network, Naomichi Matsumoto, Johi Inazawa, Magdalena Walkiewicz, Cynthia J. Tifft, Ivona Aksentijevich, Daniel L. Kastner, Pedro P. Rocha, Achim Werner

**Affiliations:** Cardiovascular and Inflammatory Disease Branch, National Human Genome Research Institute, NIH, Bethesda, MD 20892, USA; Stem Cell Biochemistry Unit, National Institute of Dental and Craniofacial Research, NIH, Bethesda, MD 20892, USA; Unit on Genome Structure and Regulation, National Institute of Child Health and Human Development, NIH, Bethesda, MD 20892, USA; Department of Molecular Cytogenetics, Medical Research Institute, Tokyo Medical and Dental University, Tokyo, Japan; Department of Human Genetics, Graduate School of Medicine, Yokohama City University Graduate School of Medicine, Japan; Office of the Clinical Director, National Human Genome Research Institute, NIH, Bethesda, MD 20892, USA; Department of Genetic Medicine, Johns Hopkins Hospital, Baltimore, MD 21287, USA; Department of Neurology, Johns Hopkins Hospital, Baltimore, MD 21287, USA; Division of Metabolic Disorders, CHOC Children’s Specialists, Orange, CA 92868, USA; Department of Pediatrics, University of California Irvine School of Medicine, Orange, CA 92868; Department of Human Genetics, Radboud University Medical Center, Nijmegen, the Netherlands; Department of General Perinatology, Hiroshima City Hiroshima Citizens Hospital, Japan; Department of Pediatrics, Hiroshima University Graduate School of Biomedical and Health Sciences, Japan; Department of Medical Genetics, Shinshu University, School of Medicine, Nagano, Japan; Bioinformatics and Scientific Programming Core, National Institute of Child Health and Human Development, NIH, Bethesda, MD 20892, USA; Undiagnosed Diseases Program, National Human Genome Research Institute, NIH, Bethesda, MD 20892; National Institute of Allergy and Infectious Disease, National Institutes of Health, Bethesda, MD 20892, USA; National Cancer Institute, NIH, Bethesda, MD 20892, USA

## Abstract

Embryonic development occurs through commitment of pluripotent stem cells to differentiation programs that require highly coordinated changes in gene expression. Chromatin remodeling of gene regulatory elements is a critical component of how such changes are achieved. While many factors controlling chromatin dynamics are known, mechanisms of how different chromatin regulators are orchestrated during development are not well understood. Here, we describe LINKED (<u>LINK</u>age-specific-deubiquitylation-deficiency-induced <u>E</u>mbryonic <u>D</u>efects) syndrome, a novel multiple congenital anomaly disorder caused by hypomorphic hemizygous missense variants in the deubiquitylase OTUD5/DUBA. Studying LINKED mutations in vitro, in mouse, and in models of neuroectodermal differentiation of human pluripotent stem cells, we uncover a novel regulatory circuit that coordinates chromatin remodeling pathways during early differentiation. We show that the K48-linkage-specific deubiquitylation activity of OTUD5 is essential for murine and human development and, if reduced, leads to aberrant cell-fate specification. OTUD5 controls differentiation through preventing the degradation of multiple chromatin regulators including ARID1A/B and HDAC2, mutation of which underlie developmental syndromes that exhibit phenotypic overlap with LINKED patients. Accordingly, loss of OTUD5 during early differentiation leads to less accessible chromatin at neural and neural crest enhancers and thus aberrant rewiring of gene expression networks. Our work identifies a novel mechanistic link between phenotypically related developmental disorders and an essential function for linkage-specific ubiquitin editing of substrate groups (i.e. chromatin remodeling complexes) during early cellfate decisions – a regulatory concept, we predict to be a general feature of embryonic development.

---

Cell-fate decisions during human development rely on signaling information encoded with ubiquitin, an essential posttranslational modifier that is reversibly attached to substrates in monomeric or polymeric forms by intricate enzymatic cascades (*1-6*). Deubiquitylases of the ovarian tumor family (OTU DUBs) are important regulators of the ubiquitin code (*7, 8*) and control crucial aspects of human physiology (*9-11*). OTU DUBs elicit their functions by hydrolyzing specific lysine linkage types within polyubiquitin to modulate the stability, activity, or interaction landscapes of their substrates (*12*). In particular, cleavage of K48-linked ubiquitin chains protects substrates from proteasomal degradation and cleavage of K63-linked ubiquitin chains limits cellular signaling. While some OTU DUBs are well characterized and have been linked to monogenetic developmental or autoinflammatory diseases (*9, 1, 13, 14*), the physiological functions and underlying mechanisms of the majority of OTU DUBs have remained largely elusive.

To identify OTU DUBs with critical roles in human development, we employed haploinsufficiency intolerance and missense constraint scores (***Fig. S1A***) (*15*). This approach revealed that *OTUD5* (also called DUBA (*16*)) is highly restricted in loss of function and missense mutations in healthy individuals, suggesting that mutations in *OTUD5* may lead to early onset disease or lethality. Through combined resources at the National Institutes of Health and external collaborations, we then queried exome sequences of a cohort of patients with developmental diseases. We identified eight male patients with novel variants in the X-linked gene *OTUD5* (***Fig. 1A***). Our index family (F1) included three brothers P1-P3, all with severe multiple congenital anomalies, all hemizygous for a maternally inherited missense mutation at p.Gly494Ser (***Fig. 1A-D***). A second novel variant, inherited *de novo,* was detected at p.Leu352Pro in patient P4, who exhibited overlapping clinical features. We further identified four additional individuals with attenuated phenotypes (***Fig. 1A,B,D***). All patients displayed global developmental delay with brain malformations, with a range of clinical characteristics including neonatal lethality and multilineage patterning defects in severe cases (***Fig. 1C-D**, **Fig. S1B, Table S1***). Although being heterozygous for the mutant allele, carrier mothers were unaffected and presented evidence of skewed X-inactivation by both methylation-specific restriction enzyme testing (***Fig. S1C***) and by RNA sequencing (see below). Consistent with previous large-scale knock out screens (*17*), we found that CRISPR-mediated knock out of OTUD5 or knock in of p.Gly494Ser or p.Leu352Pro patient alleles was lethal in mice (***Fig. S1D***). Our human and mouse genetic data thus reveal a requirement for OTUD5 for proper embryonic development.

**Fig. 1:**
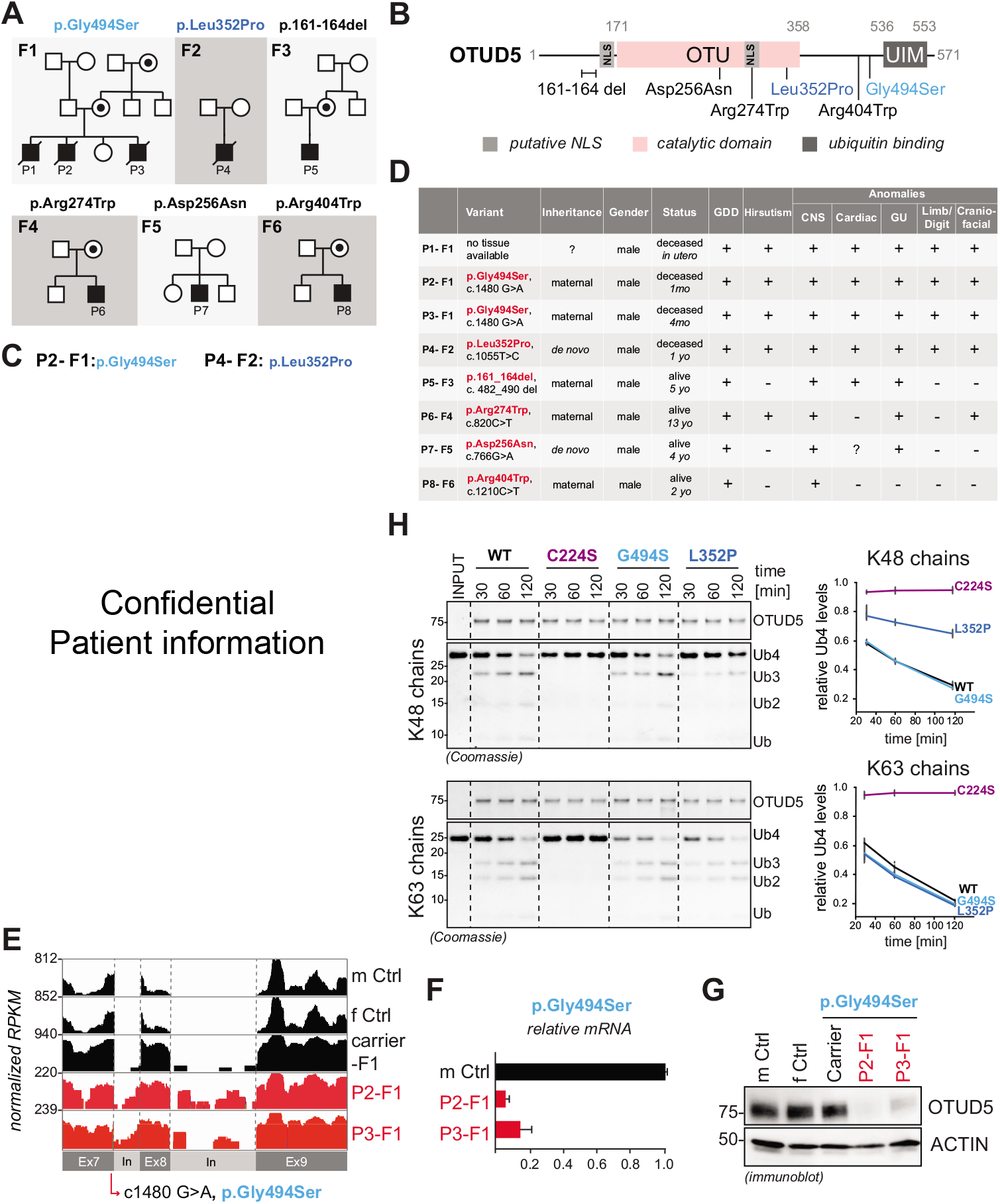
Hypomorphic hemizygous variants in OTUD5 cause developmental disease. (**A**) Genetic pedigrees of eight patients from five families with hemizygous missense mutations in OTUD5, all with overlapping phenotypes. (**B**) Domain structure of OTUD5 indicating the location of the patient mutations. Variants associated with the most severe phenotypes, p.Gly494Ser and p.Leu352Pro, are highlighted in blue colors. (**C**) Clinical photos showing craniofacial (retrognathia, midface hypoplasia, hypertelorism, low set posteriorly rotated ears, craniosynostosis) and digital anomalies (bilateral post-axial polydactyly of the hands and feet) of two patients carrying p.Gly494Ser (left panel) or p.Leu352Pro (right panel) variant of OTUD5. (**D**) Clinical table highlighting multiple congenital anomalies in patients with OTUD5 mutations. GDD= global developmental delay, CNS= Central Nervous System, GU= Genitourinary. Detailed manifestations for each category listed in Table S1. **(E**) The c.1480 G>A, p.Gly494Ser mutation is located in a 5’ splice site and leads to intron retention and reduction of OTUD5 mRNA levels as revealed by RNA sequencing of patient-derived fibroblasts. RNA sequencing reads were differentially scaled to visualize intron retention. m Ctrl = male control, f Ctrl = female control, carrier F1 = mother carrier. (**F**) The c.1480 G>A, p.Gly494Ser mutation results in a decrease in OTUD5 mRNA levels in patient-derived fibroblasts as determined by qRT-PCR. (**G**) The p.Gly494Ser mutation results in a decrease in OTUD5 protein levels as revealed by immunoblotting of lysates of patient-derived fibroblasts using indicated antibodies. (**H**) The Leu352Pro mutation specifically reduces OTUD5’s K48-ubiquitin chain cleavage activity. Wildtype ^FLAGHA^OTUD5 (WT), catalytically inactive ^FLAGHA^OTUD5 (C224S), and patient variant ^FLAGHA^OTUD5 (G494S and L352P), were purified from HEK293T cells and incubated with tetra-K48- or tetra-K63-ubiquitin chains for indicated time periods and analyzed by colloidal Coomassie-stained SDS PAGE gels. Quantification of three independent in vitro deubiquitylation experiments is shown (error bars denote s.e.m). Intensity of Ub4 band is relative to the sum of intensity of Ub3, Ub2, and Ub band.

OTUD5 is a nuclear, phospo-activated deubiquitylase that prefers cleavage of degradative K48- and non-degradative K63-ubiquitin chains over other linkage types (*8, 16, 18, 19*). To determine how the patient mutations affect OTUD5 function, we employed cell-based and biochemical assays. These studies revealed three distinct loss-of-function mechanisms. First, the OTUD5 p.Gly494Ser (c.1480 G>A) found in one of the most severe cases is located at an exon-intron splice junction and caused a reduction in *OTUD5* mRNA expression with intron retention (***Fig. 1E-F***) leading to decreased protein levels (***Fig. 1G***) likely through RNA instability. Second, p.Arg274Trp, present in a conserved putative nuclear localization sequence (***Fig. S2A***), resulted in partial mis-localization of OTUD5 to the cytoplasm (***Fig. S2A-C***). Third, with the exception of the splice-site-altering p.Gly494Ser, all patient mutations clustered around the catalytic domain of OTUD5 exhibited lower cleavage activity towards ubiquitin chains *in vitro* (***Fig. 1H, Fig. S3A-E***) without significantly affecting OTUD5’s activating phosphorylation (***Fig. S3F***). Strikingly, p.Leu352Pro, associated with a severe phenotype, specifically impaired cleavage of K48-but not K63-ubiquitin chains (***Fig. 1H, Fig. S3D-E***), highlighting an important contribution of loss of degradative chain cleavage to the disease. We thus conclude that proper levels, nuclear localization, and specifically K48-ubiquitin chain editing activity of OTUD5 are critical for its role during development.

OTUD5 had been predominantly investigated as a regulator of immune signaling (*16, 20*). However, our discovery that hypomorphic mutations in *OTUD5* cause a severe developmental disease, without any immune manifestations, gave us the unique opportunity to study the role of OTUD5, in particular its K48-deubiquitylation activity, during early cell-fate decisions of human embryogenesis. We focused on the splice-site-altering p.Gly494Ser and K48-cleavage-deficient p.Leu352Ser variants, both associated with severe phenotypes. First, we established patient-derived induced pluripotent stem cells (iPSCs) and performed teratoma formation assays. Since affected patients had craniofacial and structural brain malformations, we concentrated on ectoderm differentiation and found defects in patient cells expressing the p.Gly494Ser allele (***Fig. S4A***). Based on these observations, we subjected these iPSC lines to dual-SMAD inhibition (neural conversion), which directs differentiation towards central nervous system (CNS) precursor and neural crest cells (*21*) (***Fig. 2A***). We observed a marked upregulation of OTUD5 levels during differentiation of iPSCs of the mother carrier, suggesting a functional role for OTUD5 during this process (***Fig. 2B***). Indeed, while we observed no significant differences in the expression of pluripotency markers OCT4 and NANOG, there was a striking defect in the neural differentiation capacity when comparing iPSCs of affected patients and to carrier mother (***Fig. 2B***). This was apparent by the loss of neural crest markers, including SOX10 and SNAIL2, and the aberrant expression of CNS markers, including increases in the forebrain marker FOXG1 and decreases in neural stem cell marker PAX6, as evidenced by immunoblotting and qPCR (***Fig. 2B-C***) or at single-cell resolution by immunofluorescence (***Fig. 2D***). These results were corroborated by neural conversion experiments using hES H1 cells, in which shRNA- or siRNA-mediated depletion of OTUD5 resulted in a similar reduction of neural crest cell progeny and aberrant formation of CNS precursor cells (***Fig. S4B-E***). Importantly, these effects could be rescued by shRNA-resistant wild type OTUD5, but not by catalytically inactive C224S or K48-cleavage-deficient p.Leu352Ser OTUD5 (***Fig. 2E-F, Fig. S4F***). We therefore conclude that during embryonic development, OTUD5 regulates cell-fate decisions by specifically editing degradative K48-ubiquitin linkages on substrates and when mutated leads to a multiple congenital anomaly disease we name LINKED (<u>LINK</u>age-specific-deubiquitylation-induced <u>E</u>mbryonic <u>D</u>efects) syndrome.

**Fig. 2:**
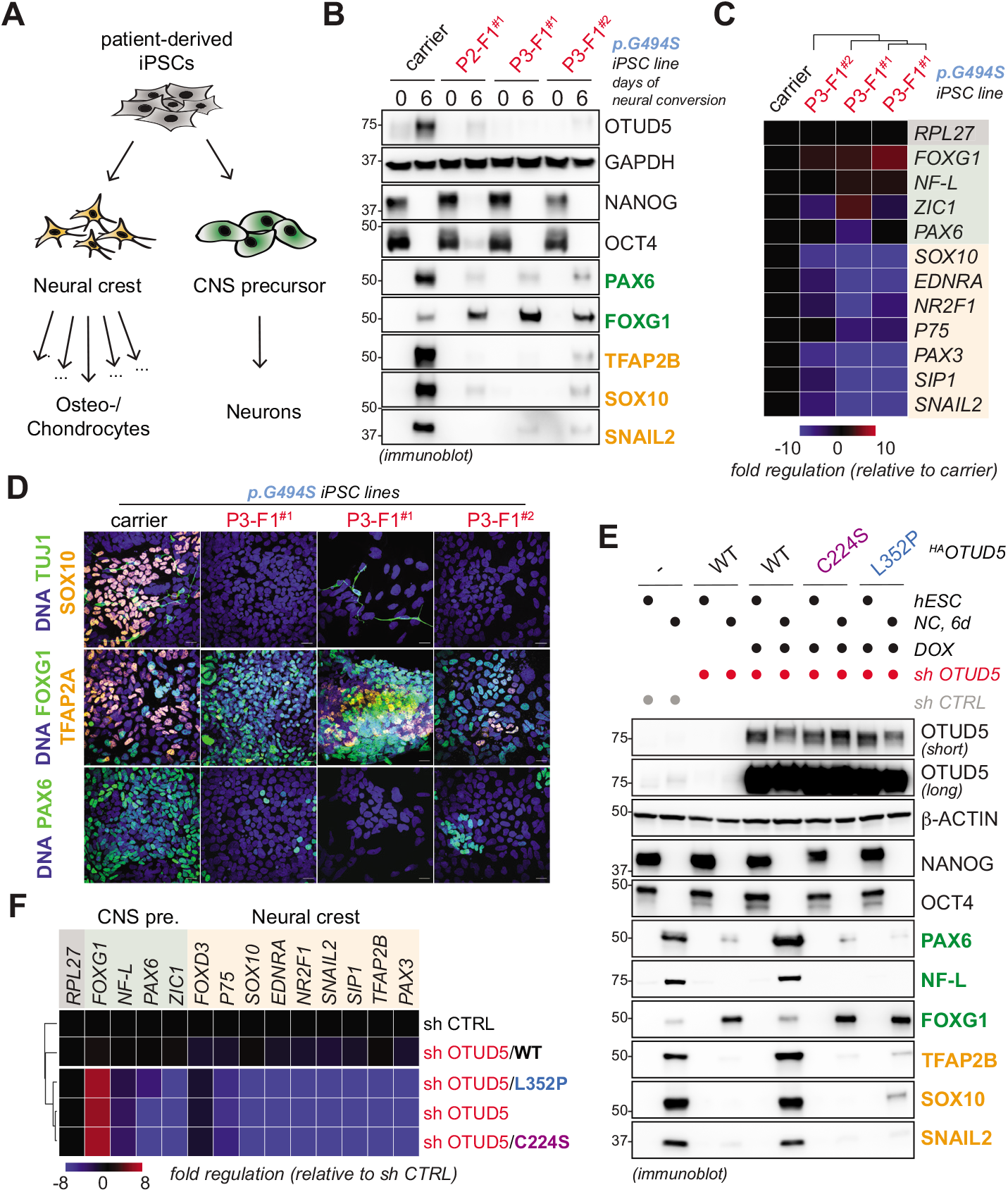
OTUD5 regulates CNS precursor and neural crest cell differentiation via its K48-ubiquitin chain specific deubiquitylation activity. (**A**) Schematic overview of neural conversion, a differentiation paradigm that directs differentiation of human pluripotent stem cells towards central nervous system (CNS) precursor cells that can further differentiate into neurons or towards neural crest stem cells, a multipotent stem cell population that can give rise to diverse cell types including craniofacial chondrocytes and osteocytes. (**B**) Reduction of OTUD5 levels causes aberrant neural conversion. iPSCs derived from OTUD5 p.Gly494Ser patients or the maternal carrier were subjected to neural conversion for 6 days. Differentiation was monitored by immunoblotting using antibodies against NANOG and OCT4 (hESC markers), PAX6 and FOXG1 (CNS precursor markers, green), TFAP2B, SNAIL2, and SOX10 (neural crest markers, orange) and GAPDH (loading control). (**C**) Reduction of OTUD5 levels causes aberrant neural conversion. iPSCs derived from the pGly494Ser patients and the mother carrier were subjected to neural conversion for 6 days and analyzed by qRT-PCR for expression of CNS precursor markers (highlighted in green) and neural crest markers (highlighted in orange). Marker expression was normalized to carrier control followed by hierarchical cluster analysis. RPL27 was used as endogenous control. (**D**) Reduction of OTUD5 levels causes aberrant neural conversion, as seen at the single cell level. Patient or mother control-derived iPSCs were subjected to neural conversion for 9d and the success of differentiation was determined by immunofluorescence microscopy using antibodies against PAX6 and FOXG1 (CNS precursor markers), TUJ1 (neuronal marker), and TFAP2A and SOX10 (neural crest markers). Scale Bar = 20μm. (**E**) K48-ubiquitin chain specific deubiquitylation activity of OTUD5 is required for proper CNS precursor and neural crest differentiation. hES H1 cells stably expressing shRNA-resistant and doxycycline-inducible wildtype (WT), catalytically inactive (C224S), or K48-chain cleavage deficient (L352P) ^HA^OTUD5 were generated. Cells were then depleted of endogenous OTUD5 using shRNA as indicated, treated with or without doxycycline (DOX), and subjected to neural conversion for 6 days. This was followed by immunoblotting using the indicated antibodies against hESC, CNS precursor, and neural crest markers. (**F**) K48-chain specific deubiquitylation activity of OTUD5 is required for proper differentiation into CNS precursor and neural crest cells. H1 hESCs reconstituted with wildtype and specific OTUD5 variants were generated and subjected to neural conversion as described above, followed by qRT-PCR analysis for expression of CNS precursor markers (highlighted in green) and neural crest markers (highlighted in orange). Marker expression was normalized to sh control followed by hierarchical cluster analysis. RPL27 was used as endogenous control.

Our results so far suggested that OTUD5 activity controls differentiation, but is less important for selfrenewal of hESCs. Therefore, we reasoned that any substrates through which OTUD5 regulates neural differentiation, should be i) more ubiquitylated in the absence of OTUD5 during neural conversion and ii) found in a physical complex with OTUD5. To identify these essential substrates, we performed a series of proteomic experiments (***Fig. 3A***). First, we employed Tandem Ubiquitin Binding Entity (TUBE)-based mass spectrometry to capture high confidence ubiquitylated proteins during neural conversion of control and OTUD5-depleted hESCs. Consistent with a role of OTUD5 during cell-fate determination, these experiments revealed that OTUD5 regulates ubiquitylation networks preferentially during neural conversion and less in self-renewing hESCs (***Fig. 3B, Table S2***). Second, we used mass spectrometry to identify proteins that bound OTUD5 during hESC differentiation (***Table S3***). Combining these two data sets, we found ~40 high probability OTUD5 substrates that were more ubiquitylated in OTUD5-depleted differentiating hESCs and were OTUD5 interactors (***Fig. S5A, Table S4***). Intriguingly, chromatin regulators were significantly enriched in these high probability substrates (***Fig. 3B, Fig. S5B***), including the BAF complex components ARID1A/B, the histone deacetylase HDAC2, the transcriptional regulator HCF1, and the ubiquitin E3 ligase UBR5, all of which had previously been shown to control neural cell-fate decisions (***Fig. S5A***) (*22-27*). We confirmed interactions with these chromatin regulators by immunoblotting at the endogenous level in hESCs (***Fig. S5C***) and demonstrated that the C-terminus of OTUD5 including the UIM motif is necessary and, with the exception of UBR5, sufficient for these binding events (***Fig. 3C***).

**Fig. 3:**
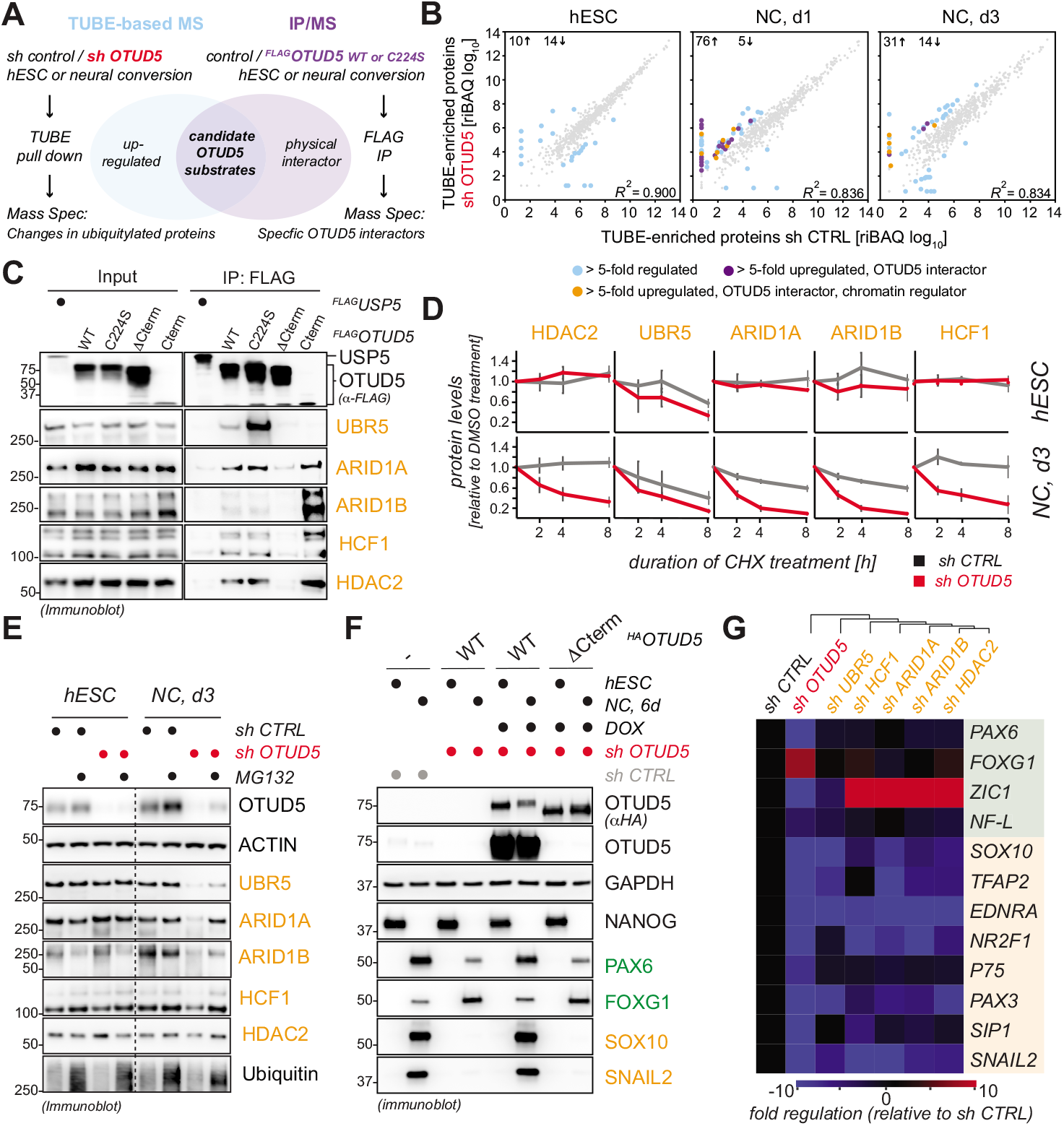
OTUD5 controls early embryonic differentiation through regulating the stability of chromatin remodelers. (**A**) Strategy used to isolate high-probability substrates of OTUD5. Two independent proteomic experiments were performed. First, control or OTUD5-depleted H1 hESCs or hESCs undergoing neural conversion for 1 or 3 days were lysed and ubiquitylated proteins were isolated by TUBE pull down followed by protein identification via mass spectrometry. Second, self-renewing or differentiating control hESCs or hESCs expressing wildtype (WT) or catalytically inactive (C224S) ^FLAG^OTUD5 were lysed and subjected to anti-FLAG immunoprecipitation followed by identification of interacting proteins via mass spectrometry. Candidate OTUD5 substrates were defined as proteins found to be more ubiquitylated upon OTUD5 depletion and identified as specific OTUD5 WT or C224S interactors. (**B**) OTUD5 preferentially controls ubiquitylation dynamics during neural conversion and many OTUD5 candidate substrates are chromatin regulators. High probability ubiquitylated proteins of control and OTUD5-depleted hESCs were identified by TUBE-based mass spectrometry as described above and relative iBAQ values were plotted against each other for each differentiation state (hESC, NC day 1, NC day 3). More than 5-fold regulated proteins are highlighted in blue and total number of upregulated or downregulated proteins are indicated in the upper left corner of the diagram for each differentiation state. More than 5-fold upregulated proteins also found in ^FLAG^OTUD5 IPs (i.e. candidate OTUD5 substrates) are highlighted in violet. Note that candidate OTUD5 substrates are found specifically during differentiation, but not in hESCs. Candidate OTUD5 substrates are significantly enriched in chromatin binding proteins (highlighted in orange) as determined by GO analysis. (**C**) OTUD5 interacts with chromatin regulators via its C-terminus. HEK293T cells transiently expressing ^FLAG^OTUD5 wildtype and indicated mutants were lysed and lysates were subjected to anti-FLAG immunoprecipitation followed by SDS PAGE and immunoblot analysis using indicated antibodies. (**D**) OTUD5 stabilizes chromatin regulators in differentiating, but not self-renewing hESCs. Control or OTUD5-depleted hESCs or hESC subjected to neural conversion for 3 days were treated with cycloheximide for indicated time periods and protein stability of HDAC2, UBR5, ARID1A/B, and HCFC1 was determined by immunoblotting. Quantification of three biological replicates is shown (error bars denote s.d., chromatin regulator levels were normalized relative to actin levels and 0h time point set to 1). (**E**) OTUD5 protects chromatin regulators from proteasomal degradation in differentiating, but not self-renewing hESCs. Control or OTUD5-depleted hESCs or hESC subjected to neural conversion for 3 days were treated with the proteasome inhibitor MG132 for 4h followed by immunoblotting with indicated antibodies. (**F**) The chromatin regulator binding-deficient OTUD5^DCterm^ mutant does not support neural conversion. hES H1 cells stably expressing shRNA-resistant and doxycycline-inducible wildtype (WT) or chromatin regulator binding-deficient (DCterm) ^FLAGHA^OTUD5. Cells were then depleted of endogenous OTUD5 using shRNA as indicated, treated with or without doxycycline (DOX), and subjected to neural conversion for 6 days. Successful differentiation was monitored by immunoblotting using the indicated antibodies against hESC, CNS precursor, and neural crest markers. Note that anti-OTUD5 antibodies were raised against the C-terminus of OTUD5 and thus do not recognize OTUD5^DCterm^. (**G**) Individual depletion of chromatin regulators partially phenocopies the aberrant neural conversion program observed upon OTUD5 reduction. hES H1 cells were depleted of endogenous OTUD5 or indicated chromatin regulators using stably expressed shRNAs and subjected to neural conversion for 6 days and analyzed by qRT-PCR for expression of CNS precursor markers (highlighted in green) and neural crest markers (highlighted in orange). Marker expression was normalized to sh control followed by hierarchical cluster analysis. RPL27 was used as endogenous control.

OTUD5 controls ectodermal differentiation via its K48-cleavage activity (***Fig. 2***) and K48-ubiquitin chains are a common targeting signal for proteasomal degradation (*1*). Thus, if the interacting chromatin regulators were substrates of OTUD5, we expected to see changes in their stability in the absence of OTUD5. To test this hypothesis, we performed cycloheximide chases in self-renewing and differentiating control or OTUD5-depleted hESCs. Indeed, while reduction of OTUD5 had no significant impact on the half-life of the chromatin regulators in self-renewing hESCs, it dramatically decreased their stability during neural conversion (***Fig. 3D, Fig. S5D***), a process that was dependent upon the proteasome (***Fig. 3E***). To investigate whether OTUD5 regulates neuroectodermal differentiation through targeting these substrates, we next performed rescue experiments using a version of OTUD5 that lacked its C-terminus (OTUD5^ΔCterm^) and thus was deficient in chromatin regulator binding (***Fig. 3C***), yet retained K48-specific deubiquitylation activity (***Fig. S6A***). Intriguingly, this separation-of-function mutant failed to support neural crest differentiation and showed aberrant CNS precursor formation (***Fig. 3F, Fig. 6B***). In line with this observation, individual depletion of these chromatin regulators resulted in a similar, yet less pronounced dysregulation of the neural conversion program as the one observed upon loss of OTUD5 (***Fig. 3G, Fig. S6C***). Taken together, these results suggest that during early differentiation, OTUD5 edits K48 ubiquitin chains deposited on a subset of chromatin regulators to prevent their proteasomal degradation, thus coordinating their function in chromatin remodeling events required for neural cellfate commitment. Consistent with this notion, loss-of-function mutations in these chromatin regulators lead to Coffin Siris (ARID1A/B (*27*)) and Cornelia de Lange syndromes (HDAC2 (*26*)), broad spectrum developmental diseases that exhibit phenotypic overlap with LINKED syndrome patients (***Fig. 1, Fig. S7***).

Given that OTUD5 controls the stability of several chromatin regulators such as ARID1A and ARID1B during early ectodermal cell-fate commitment, we hypothesized that the observed clinical and differentiation phenotypes underlie impaired chromatin dynamics. In line with this, ATAC-seq revealed that OTUD5 depletion caused loss of accessible chromatin at early stages of neural conversion, while accessibility was largely unaffected in self-renewing hESCs (***Fig. 4A-B, Fig. S8A-B***). This was accompanied by modest transcriptional changes (***Fig. S8C-D***), suggesting that differences in ATAC signal are not a consequence of failed differentiation, but rather its initiating cause. Genes associated with chromatin regions exhibiting reduced accessibility at day 3 of neural conversion were enriched for GO terms involved in specification of CNS precursor and neural crest cells (***Fig 4C***). Strikingly, more than half of the regions displaying lower accessibility were located at enhancers (***Fig 4D***), and were enriched for neural and neural crest fate-promoting transcription factor motifs such as OTX2, SOX2, SOX3, SOX9 and SOX10 (***Fig. S9***). Further supporting the idea that OTUD5 regulates cell-fate commitment through stabilizing chromatin remodelers, we observed that OTUD5-regulated neural enhancers were often bound by SMARCA4(*28*), a component of the BAF complex recruited by ARID1A/B (*29*), in neural progenitor cells (***Fig. 4B***). Among the ~600 genes associated with less accessible enhancers was the neuroectoderm cell-fate promoting transcription factor *PAX6* and the neural crest and neuronal specification factor *SEMA3A*. These two cell-fate regulators exhibited reduced mRNA expression upon OTUD5 depletion at these early stages of differentiation (***Fig. 4E***). Thus, OTUD5 controls neural cell-fate commitment by regulating chromatin accessibility at neural- and neural crest-specific enhancers to enable activation of transcriptional networks that drive the differentiation program.

**Fig. 4:**
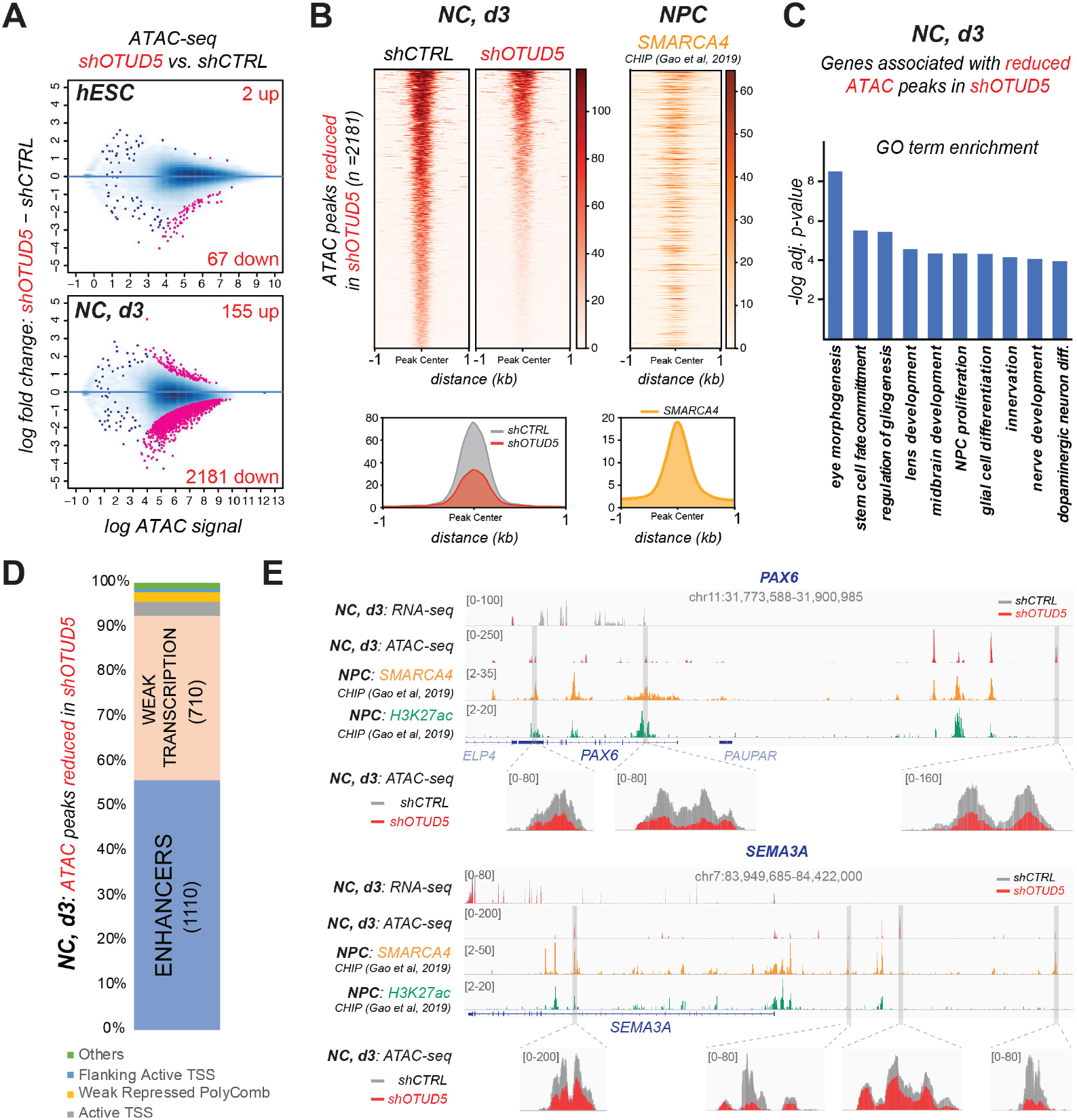
OTUD5 is required for chromatin remodeling at enhancers driving neural and neural crest differentiation. (**A**) Loss of OTUD5 leads to changes in chromatin accessibility specifically during differentiation. Changes in ATAC-seq signal depicted as log two-fold change and signal intensity in hES H1 cells (hESC) and cells subjected to neural conversion for 3 days (NC, d3). Signal at stringently identified peaks (IDR 0.05) was compared between control and OTUD5-depleted cells. Pink dots represent peaks with statistically significant enrichment differences (adj pvalue < 0.0001) between the two conditions. There is a small number of differentially enriched ATAC-seq peaks in self-renewing hES H1 cells (top MA plot), while during neural conversion the majority of changes occur at ATAC peaks that lose accessibility upon OTUD5 depletion (bottom MA plot). Numbers of statistically significant peaks gaining (up) or losing (down) accessibility upon OTUD5 depletion are indicated. (**B**) Pooled ATAC-seq signal from three independent replicates for each condition (shCTRL and shOTUD5) was plotted at the subset of peaks that lost chromatin accessibility upon OTUD5 depletion (total 2181). The loss of accessibility associated with OTUD5 depletion at NC, d3 is depicted as a heatmap (top panel) where averaged ATAC-signal from control or OTUD5-depleted cells is plotted. The average profile of ATAC-signal in shControl and shOTUD5 cells (bottom panel) also highlights the overall reduction in accessibility concomitant with a loss of OTUD5. Regions where chromatin accessibility is lost upon OTUD5 depletion are also commonly bound by SMARCA4, a component of the BAF complex recruited to chromatin by the OTUD5 substrates ARID1A/B, as profiled by ChIP-seq in neuronal precursor cells. by ChIP-seq in neuronal precursor cells. Bottom right plot shows that SMARCA4 peaks are centered at the differentially enriched ATAC-seq peaks. (**C**) OTUD5 is required for chromatin remodeling at genes promoting neural differentiation. ATAC-seq peaks significantly regulated by OTUD5 at day 3 of neural conversion were associated with genes using GREAT analysis followed by GO-term analysis. (**D**) OTUD5 is predominantly required for chromatin remodeling at enhancers. ATAC-seq regions with less enrichment in OTUD5-depleted differentiating cells were classified using ChromHMM genome functional annotation of H1-derived neuronal precursor cells. (**E**) Browser snapshots of the PAX6 (top) and SEMA3A (bottom) loci at day 3 of neural conversion, comparing changes in transcription (profiled by RNA-seq) and chromatin accessibility (profiled by ATAC-seq) induced by OTUD5 depletion (red) as compared to control (grey). Bottom two tracks show enrichment of H3K27ac (a histone posttranslational modification associated with active enhancers) and SMARCA4 (a component of the BAF complex recruited to chromatin by the OTUD5 substrates ARID1A/B). The reduction in ATAC-seq signal at some of the ATAC-seq peaks is associated with strong transcriptional downregulation for PAX6 and a modest, albeit significant, impact on SEMA3A.

We here discover LINKED syndrome, a novel multiple congenital anomaly disorder cause by variants in OTUD5. By mechanistically studying these mutations, we identify an important role for linkage-specific ubiquitin chain editing during embryonic development and aberrant degradation of chromatin regulators as a major disease mechanism underlying LINKED syndrome (***Fig. 5***). The K48-chain specific cleavage activity of OTUD5 is required to stabilize the levels of several chromatin remodelers during early stages of differentiation. We propose that in this manner, OTUD5 coordinates chromatin remodeling networks that promote the accessibility of enhancers to ensure transcriptional changes required for ectodermal cellfate commitment.

**Fig. 5:**
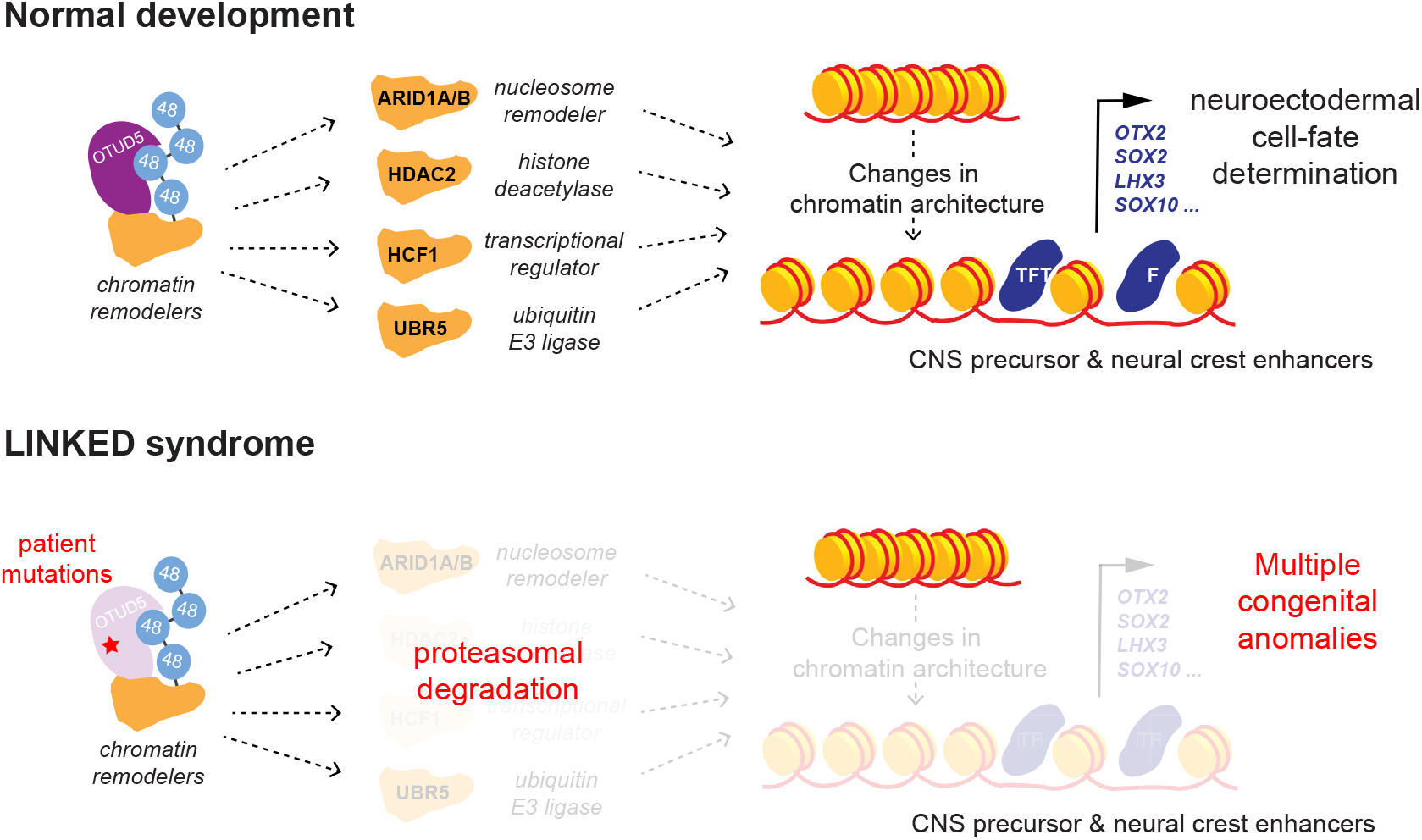
OTUD5 controls developmental chromatin dynamics and when mutated leads to LINKED syndrome. Model of how linkage-specific ubiquitin chain editing by OTUD5 controls development and is misregulated in disease. During normal early embryogenesis, OTUD5 employs its K48-linkage specific deubiquitylation activity to target and stabilize several key chromatin regulators to coordinate chromatin remodeling events at CNS precursor and neural crest enhancers. This allows binding of lineage-promoting transcription factors to drive transcriptional networks required for neuroectodermal cellfate commitment. Hypomorphic patient mutations in OTUD5 result in dysregulation of this pathway and lead to a novel multiple congenital anomaly disorder we name LINKED syndrome.

Our work has important implications for our understanding of ubiquitin-dependent signaling during embryonic development. Rather than targeting one particular protein, OTUD5 acts by cleaving K48-ubiquitin chains off a group of functionally related substrates, i.e. chromatin remodelers, to drive differentiation. Together with genetic studies connecting *OTUD7A* and *OTUD6B* mutations to neurodevelopmental diseases (*30-32*), our data therefore suggest that signaling through linkage-specific ubiquitin editing of substrate groups by OTU DUBs will be a common mechanism utilized to coordinate embryonic differentiation pathways.

Coffin Siris and Cornelia de Lange syndrome are multiple congenital anomaly disorders that, while exhibiting considerable phenotypic variability and allelic as well as locus heterogeneity, share disease manifestations. Here, we describe LINKED syndrome, which exhibits aspects both distinct from, and in common with, these two syndromes (***Fig. S7***). OTUD5 targets ARID1A/B and HDAC2, proteins that are mutated in Coffin Siris and Cornelia de Lange syndrome, respectively. Our findings thus link these proteins to a common pathway, thereby providing a molecular framework to explain the clinically overlapping features of these genetic disorders. In addition, ARID1A/B and HDAC2 are members of multi-subunit chromatin regulatory complexes that are frequently mutated in human cancers (*33, 34*). Given the rising interest in DUBs as drug targets (*35*), OTUD5 might therefore be an attractive candidate for therapeutic intervention. Determining the effects of OTUD5 loss on chromatin regulator complex stoichiometries during embryonic and cancer development and identification of cognate E3 ligases will hence be important fields of future investigation.

## Material and Methods

### Human subjects

Written informed consent was obtained from all individuals or family member legal representatives prior to exome sequencing. Consent was obtained for publication of photographs prior to inclusion in the study. Family 1, individuals 1-3, were counseled regarding the possible outcomes of exome sequencing and signed a consent form for research-based exome sequencing through the Johns Hopkins Hospital and the National Institutes of Health, which was approved by the National Institutes of Health Institutional Review Board (IRB). The rest of the participants were recruited through GeneMatcher (36). Individual 4 was consented for clinical exome sequencing through Children’s Hospital of Orange County. Individual 5 was consented for clinical exome sequencing through Johns Hopkins Hospital. Individual 6 was consented for clinical and/ or research-based exome sequencing through Tokyo Medical and Dental University. Individual 7 was consented for clinical and/or research-based exome sequencing through Radboud University, Nijmegen, Netherlands. Individual 8 was consented for clinical and/or research-based exome sequencing through Yokohama City University, Japan. Ethics oversight for patients was provided by the institutions that consented the respective patients.

### Exome and Sanger sequencing

Whole-exome sequencing and data analysis were performed as previously described. New candidate variants were filtered to remove those present in the ExAC, 1000 Genomes Project, dbSNP, NHLBI GO Exome Sequencing Project and ClinSeq databases and an in-house database with over 1200 exomes, and variants were selected on the basis of autosomal recessive or X-linked recessive inheritance. For individual 3, trio-based exome sequencing was performed on genomic DNA isolated from amniocytes through Johns Hopkins Hospital and the National Institutes of Health. Novel variant in OTUD5 was identified using standard bioinformatics analysis. Sanger sequencing confirmed the presence of the OTUD5 variants in the trio and their absence in unaffected family members. Individual 4 had standard exome sequencing performed through Children’s Hospital of Orange County and a novel variant in OTUD5 was identified and Sanger confirmed. Individual 5 was consented for clinical exome sequencing through Johns Hopkins Hospital. Individual 6 was consented for clinical and/or research-based exome sequencing through Radboud University, Nijmegen, Netherlands. Individual 7 was consented for clinical and/or research-based exome sequencing through Shinshu University, Nagano, Japan. All variants were confirmed using Sanger sequencing. All patients reported have no known definitive pathogenic variants identified in other genes causative for multiple congenital anomalies and developmental delay. All patients provided informed consent for exome sequencing and identifiable photographs. The approved study protocol for this work includes 94-HG-0105 and the UDN from NHGRI.

### Fibroblast/Induced pluripotent cell lines

Dermal fibroblast cells derived from OTUD5 patients or unrelated healthy donors were grown in DMEM (Life Technologies) supplemented with 10% FCS (Gemini Bio-Products) and 1× antibiotics (Life Technologies). Induced pluripotent stem cells (iPS) were generated from individuals 2 and 3, and their unaffected carrier mother using a 3 week sendai virus protocol previously described (37). Reprogramming efficiency was measured using both FACS analysis for pluripotency markers and teratoma formation in nude mice. Multiple clones were generated for each iPS line and were used in specific experiments. Teratoma formation assays were performed on each clone, in quadruplicate, using bilateral gastrocnemius muscle injection in each mouse as described elsewhere (38). Slides were generated and stained using Hematoxylin and eosin stain and quantified for ectodermal components using Adobe Illustrator.

### Determination of skewed X-inactivation

The methylation status of the human androgen receptor (AR) gene at Xq12 was assessed to infer X chromosome inactivation in the heterozygous mother carrying the OTUD5 p.R274W mutation. 100 ng of DNA isolated from peripheral blood was digested with the methylation-sensitive HpaII enzyme (New England Biolabs, Ipswich, MA, USA), as originally described (39). Digested and undigested samples were then amplified by PCR with primers and protocol as previously described (40). The PCR products were separated by capillary electrophoresis on an ABI 3730xl DNA Analyzer (Applied Biosystems) with the GeneScan 500 LIZ size standard (Applied Biosystems). Fragment analysis was performed with GeneMapper software (Applied Biosystems).

### Mouse studies

Transgenic mice (C57BL/6J) were generated using CRISPR-Cas9 injection and electroporation after isolation of early embryos (41). For generating Otud5 knock-in mutations we used a gRNA p.Gly494Ser 5’-CACCCTGTGCACCAGGTCAG-3’ and for p.Leu352Pro we used gRNA 5’-CCCCTGGCTTAAATGACGGT-3’ with repair templates including the specific missense mutation. After failing to identify viable pups with either indels or specific knock-in mutations, we began isolating day 12.5 embryos by microsurgery, imaging the embryo, and genotyping a small portion of the tail. As a control for injections, we either used saline injection or a non-essential gene used in the lab (Tbx21) 5’-CCCACTGTGCCCTACTACCG-3’. All mouse work included was approved by the National Human Genome Research Institute animal protocol and ethically overseen by OLAW (Office of Laboratory Animal Welfare).

### Plasmids, shRNAs, siRNAs

pDEST-FLAG-HA-USP5 and pDEST-FLAG-HA-OTUD5 were gifts from Wade Harper (Addgene plasmids # 22590 and # 22610(42). OTUD5 patient mutations (G494S, L352P, 161-164del, R274W, and D265N) point mutations (C224S), truncation mutations (DCterm = OTUD5 1-534, Cterm = OTUD5 534-571), and wobble mutations to make constructs resistant to shOTUD5#5 were introduced in this vector using the Q5 site-directed mutagenesis kit (E0554, NEB) following the manufacturer’s instructions. For expression in human embryonic stem cells, OTUD5 variants were cloned into pENTR1A or pENTR233 and recombined into pINDUCER20 (43). pLKO1-Puro Mission shRNA constructs targeting OTUD5 (#2: TRCN00001 22275, #5: TRCN0000233196), ARID1A (TRCN 0000059092), ARID1B (TRCN00000420576), UBR5 (TRCN0000003411), HDAC2 (TRCN00000004819), HCFC1 (TRCN00000001625) were purchased from Sigma. siRNA pools were purchase from Santa Cruz Biotechnology and Thermo Scientific.

### Antibodies

The following antibodies were commercially purchased for immunoblotting and immunofluorescence microscopy. Anit-OTUD5 (#20087S, clone D8Y2U, Cell Signaling, 1:1000 in IB), anti-PAX6 (#60433, clone D3A9V, Cell Signaling, 1:1000 in IB, 1:200 in IF), anti-TFAP2 (#2509, Cell Signaling, 1:1000 in IB, 1:200 in IF), anti-FOXG1 (ab18259, abcam, 1:1000 in IB, 1:100 in IF), anti-ARID1A (#12354, clone D2A8U, Cell Signaling, 1:3000 in IB), anti-ARID1B (#65747, clone E1U7D, Cell Signaling, 1:1000 in IB), anti-UBR5 (#65344, clone D6O8Z, Cell Signaling, 1:1000 in IB), anti-HDAC2 (#5113, clone 3F3, Cell Signaling, 1:3000 in IB) anti-HCFC1 (#50708, Cell Signaling, 1:1000 in IB), anti-Actin (#8691001, MP Biomedical, 1:10,000 in IB), anti-SOX10 (#89356, Cell Signaling, 1:1000 in IB, 1:100 in IF), anti-TUJ1 (#5568, clone D71G9, Cell Signaling, 1:1000 in IB, 1:200 in IF), anti-NANOG (#4903, clone D73G4, Cell Signaling, 1:1,000 in IB), anti-OCT4 (ac-8628, Santa Cruz, 1:1,000 in IB), anti-OCT4 (#75463, clone D7O5G, Cell Signaling, 1:1000 in IB), anti-SNAIL2 (#9585, clone C19G7, Cell Signaling, 1:500 in IB), anti-GAPDH (#5174, clone D16H11, Cell Signaling, 1:10,000), anti-HA (clone C29F4; Cell Signaling, 1:3,000 in IB and 1: 200 in IF), anti-Flag (F1804, clone M2, Sigma, 1:2,000). Anti-pOTUD5^Ser177^ antibodies were gift from Genetech and were described previously(19).

### Mammalian cell culture and transfections

Human embryonic kidney (HEK) 293T cells (ATTC) were maintained in DMEM with 10% fetal bovine serum. Plasmid transfections of HEK 293T cells were carried out using PEI. siRNA transfections were carried out with Lipofectamine RNAiMAX (Invitrogen) according to manufacturer’s instructions using 10 nM for each siRNA. Cells were routinely tested for mycoplasma using the MycoAlert Mycoplasma Detection Kit from Lonza (LT07-118).

### Pluripotent stem cell culture, lentiviral infections, and neural conversion

Human embryonic stem (hES) H1 cells (WA01, WiCell) were maintained under feeder free conditions on Matrigel-coated plates (#354277, BD Biosiences) in mTeSR^™^1, (#05871/05852, StemCell Technologies Inc.) and were routinely passaged with collagenase (#07909, StemCell Technologies Inc.). Cells were routinely tested for mycoplasma using the MycoAlert Mycoplasma Detection Kit from Lonza (LT07-118). Lentiviruses were produced in 293T cells by cotransfection of lentiviral constructs with packaging plasmids (Addgene) for 48–72 hr. Transduction was carried out by infecting 2x10^5^ hES H1 cells per well of a 6-well plate with lentiviruses in the presence of 6 μg/ml Polybrene (Sigma) and 10 μM Y-27632 ROCK inhibitor. For transduction of lentiviruses carrying ectopic expression vectors, cells were centrifuged at 1,000g at 30C for 90min. Media was replaced with 2mL mTESR1 containing 10 μM Y-27632 ROCK inhibitor. After 4-6d of selection with appropriate antibiotic (1 μg/ml puromycin for pLKO1-puro-shRNA constructs, 200 μg/ml G418 for pINDUCER20 constructs), hES H1 cells were analyzed and used in differentiation experiments. Neural induction of hES H1 cells expressing different shRNA constructs was performed using STEMdiff^™^ Neural Induction Medium (#05831, StemCell Technologies Inc.) in combination with a monolayer culture method according to the manufacturer’s technical bulletin (#28044) and as previously described (21). In brief, single cell suspensions were prepared by treatment of hES cells with accutase and 1.5 − 2.0 x 10^6^ cells were seeded per well of a 6-well plate in 4mL STEMdiff^™^ Neural Induction Medium supplemented with 10 μM Y-27632 ROCK inhibitor. Neural induction was performed for indicated time periods with daily medium change.

### hES H1 rescue experiments

To rescue OTUD5-dependent phenotypes, hES H1 cells were stably transduced with pINDUCER-^FLAGHA^OTUD5 constructs (WT, C224S, L352P or ΔC-term, containing wobble mutations that render them resistant to shOTUD5#5). Cells were selected and maintained with 200 μg/ml G418 for 4-5d. Cells were then transduced with control shRNAs or shRNAs targeting OTUD5 (shOTUD5#5) and selected and maintained with 1 μg/ml puromycin. For the rescue experiments, these cell lines were then treated in absence or presence of 1 μg/ml doxycycline and subjected to neural conversion for indicated time periods. Cells were harvested for immunoblotting and RNA extraction.

### Proteasome inhibitor treatment

To inhibit proteasome-mediated degradation of proteins, hES H1 cells and hES H1 cells undergoing neural conversion for three days were treated with the proteasome inhibitor MG132 at a concentration of 10 μM for 4h. After treatment, the cells were harvested by scraping in 1xPBS and centrifuged at 300g for 5 minutes. Cells were lysed in 2x urea sample buffer (150mM Tris pH 6.5, 6M urea, 6% SDS, 25% glycerol and a few grains of bromophenol blue) followed by immunoblotting with indicated antibodies.

### Cycloheximide (CHX) chase assays

For cycloheximide chase assays, control or OTUD5-depleted hES H1 cells and cells that had undergone neural induction for 3 d were treated with 40 mg/mL CHX for 2, 4, and 8 h. Cells were lysed in 2x urea sample buffer (150mM Tris pH 6.5, 6M urea, 6% SDS, 25% glycerol and a few grains of bromophenol blue), sonicated, and were analysed by immunoblotting. For quantification, immunoblot signals for respective proteins were quantified using ImageJ (NIH, http://rsbweb.nih.gov/ij/) and normalized to GAPDH or β-ACTIN.

### Quantitative real-time PCR (qRT-PCR) analysis

For qRT–PCR analysis, total RNA was extracted and purified from cells using the NucleoSpin RNA kit (#740955, Macherey Nagel) and transcribed into cDNA using the SuperScript^™^ IV First-Strand Synthesis System (#18091050, ThermoFisher Scientific). Gene expression was quantified by PowerUp SYBR Green qPCR (#A25741, ThermoFisher Scientific) on a CFX96 Real-Time System (Bio-Rad). Nonspecific signals caused by primer dimers were excluded by dissociation curve analysis and use of non-template controls. Loaded cDNA was normalized using RPL27 as an endogenous control. Gene-specific primers for qRT–PCR were designed by using NCBI Primer-Blast. Primer sequences can be found in Supplementary Table 5.

### Cluster analysis

mRNA abundance was measured by RT-qPCR for different conditions. The datasets were plotted as a heatmap in Python using the Seaborn library. Hierarchical clustering of samples was performed using the Bray-Curtis method with average linkage.

### Immunoprecipitations

HEK 293T cells were transiently transfected with wildtype ^FLAGHA^OTUD5 or indicated variants and incubated for 48 hours at 37°C with 5% CO_2_. Cells were harvested by scraping in 1xPBS and centrifuged at 300xg for 5 minutes. The cell pellets were either stored at -80°C or directly used for immunoprecipitation experiment. To detect OTUD5 interaction partners, HEK 293T expressing indicated ^FLAGHA^OTUD5 variants (3x15 cm dishes per condition) were lysed in two pellet volumes of ice-cold lysis buffer (20 mM HEPES pH 7.3 containing 110 mM potassium acetate, 2 mM magnesium acetate, 1 mM EGTA, 2 mM EDTA, 0.1% NP-40, 1x protease inhibitors (Roche), 1x Phos-Stop (Roche), and 2 mM phenanthroline. Cells were sonicated and the lysates were cleared by centrifugation at 20,000xg for 25min. To remove residual lipids, the supernatant was filtered through 0.22 um filter (Millex-GV). Subsequently, the lysates were quantified using Pierce 660nm reagent (Thermo, #22660) and an equal amount of lysates were incubated with ANTI-FLAG-M2 agarose (Sigma) for 2h at 4 °C. Beads were then washed three times with lysis buffer and eluted in lysis buffer supplemented with 0.5mg/mL 3xFLAG peptide (Sigma). Eluted proteins were precipitated by adding 20% TCA followed by overnight incubation on ice. Protein pellets were washed three times with ice-cold 90% acetone in 0.01 M HCl, air dried, and solubilized with 2x urea sample buffer followed by immunoblot analysis.

To detect endogenous OTUD5 interactions, anti-OTUD5 immunoprecipitations were performed from hES H1 cells (5 x 15cm dishes per condition) and lysates were prepared as described above. After incubation with OTUD5 antibodies or control antibodies (rIgGs) at 4C for 1h, Protein A beads (Roche) were added for 2h. After washing with lysis buffer, bound proteins were eluted with 2x urea sample buffer, followed by SDS page and immunoblotting using the indicated antibodies. For in vitro deubiquitylation assays, OTUD5 and indicated variants were purified from HEK 293T cells (1X15 cm dishes per condition). Lysates were prepared and subjected to anti-FLAG immunoprecipitation as described above. Beads were washed twice with lysis buffer containing 1M NaCl, three times with lysis buffer without NaCl, and OTUD5 was eluted from the beads with lysis buffer containing 10mM DTT and 0.5mg/mL 3xFLAG peptide. For mass spectrometry analysis, self-renewing or differentiating (neural conversion, 3d) hES H1 cells or hES H1 cells expressing wild type (WT) or catalytically inactive (C224S) ^FLAGHA^OTUD5 were lysed and subjected to anti-FLAG immunoprecipitation as described above (5 x15 cm dishes per condition). FLAG immunoprecipitates were further processed for multi-dimensional protein identification technology (MUDPIT) mass spectrometry as described below.

### In vitro deubiquitylation assays

For in vitro deubiquitylation reactions, equal amounts of wild type OTUD5 or indicated OTUD5 mutants (purified as described above) were incubated with 0.5 μM K48- or K63-tetra ubiquitin chains (Boston Biochem) in cleavage buffer (110 mM potassium acetate, 2 mM magnesium acetate, 1 mM EGTA, 20 mM HEPES (pH 7.3), 0.1%NP-40, 2 mM EDTA and 10 mM DTT) at 30°C for different time periods (30’, 60’, and 120’). Reactions were stopped by addition of equal amounts of 2x urea sample buffer.

### Mass spectrometry to identify OTUD5 interactors

For mass spectrometry analysis, flag-immunoprecipitates were prepared from self-renewing and differentiating hESCs as described above and precipitated with 20% Trichloroacetic acid (TCA, Fisher) overnight. Proteins were resolubilized and denatured in 8M Urea (Fisher), 100 mM Tris (pH 8.5), followed by reduction with 5 mM TCEP (Sigma), alkylation with 10 mM iodoacetamide (Sigma), and overnight digestion with trypsin (0.5 mg/ml, Fisher). Samples were analyzed by MUDPIT mass spectrometry by the Vincent J. Coates Proteomics/Mass Spectrometry Laboratory at UC Berkeley. High confidence interactors of OTUD5 were defined as nuclear proteins only found in ^FLAGHA^OTUD5^WT/C224S^ and not in control immunoprecipitates.

### TUBE-based mass spectrometry

Tandem Ubiquitin Binding entities (TUBE)-based mass spectrometry was performed by LifeSensors. The company provided the number of peptides detected, raw intensities, as well as calculated iBAQ values. The data was further processed and analyzed using a Python script. Subsequent figures were produced using Matplotlib and Seaborn libraries. First, all the runs were normalized to each other using a sum normalization method which we called the relative iBAQ values. This was done by taking the iBAQ values of each sample/run and dividing by the sum total of all iBAQ values for each sample/run.

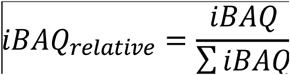

We then used the minimum detection limit to fill any missing data. All zero/NaN values were replaced by the minimum relative iBAQ value in each individual sample. Afterwards, the data was log transformed before plotting. However, the sum normalized values give a mixture of positive and negative numbers after log transformation, so all the data was multiplied by

1.0x10^7^ in order to bring the smallest values above 1 in all the samples. The log transformed data was plotted on scatter plots using Matplotlib and Seaborn. To enrich for proteins regulated in their ubiquitylation status in an OTUD5-dependent manner, we filtered for nuclear proteins with at least 13 unique peptides and identified in at least 4 of the 6 groups (depicted in grey in Figure 3b) and in addition were found more than 5-fold regulated upon OTUD5 depletion in TUBE pull downs from hESCs or hESC undergoing neural conversion for 1d or 3d (depicted in light blue in Figure 3b).

High confidence OTUD5 substrate identification:

To identify high confidence OTUD5 substrates, we filtered for proteins that we found more than 5-fold upregulated in the TUBE-based mass spectrometry and to be physically interacting with OTUD5 in our MUDPIT mass spectrometry experiments. The list of these proteins was subjected to GO term analysis.

### Immunofluorescence microscopy

For immunofluorescence analysis, hES H1 cells were seeded on Matrigel-coated coverslips using accutase, fixed with 4% formaldehyde for 20 min, permeabilized with 0.5% Triton for 10 min, and stained with indicated antibodies and/or Hoechst 33342. Images were taken using a Nikon A1R+ MP microscope and processed using ImageJ.

### ATAC-seq library preparation and downstream processing

ATAC-seq was performed using the OMNI-ATAC protocol as previously described(44). Briefly, cells were dissociated using Accutase, and 50,000 cells were subjected to the tagmentation reaction. Cells were first washed in resuspension buffer (10 mM Tris-HCl pH 8.0, 10 mM NaCl, and 3 mM MgCl2 in water), following which nuclei were isolated in 1 ml lysis buffer (10 mM Tris-HCl pH 8.0, 10 mM NaCl, 3 mM MgCl2, 0.1% NP-40, 0.1% Tween-20, and 0.01% Digitonin in water) on ice for three minutes. Nuclei were rinsed once in wash buffer (10 mM Tris-HCl pH 8.0, 10 mM NaCl, 3 mM MgCl2, and 0.1% Tween-20) and tagmentation was carried out using 2.5ul Tn5 transposase (Illumina 15027865) for 30 mins. Following tagmentation, DNA was purified using the Zymo DNA Clean and Concentrator kit. Libraries were prepared by PCR using Q5 High Fidelity DNA polymerase (NEB) polymerase and using primers carrying Illumina Nextera i7 barcodes. First, gap filling was performed at 72°C for 5mins followed by five cycles of 98°C, 20secs, 63°C, 30secs, and 72^0^C 1min. After initial amplification, tubes were held on ice, while quantitative PCR was run on 1ml of the pre-amplified library to determine additional number of cycles needed. Libraries were sequenced on HiSeq2500 using PE50. Raw reads were processed using the ENCODE pipeline (encodeproject.org/atac-seq/) (45). Differentially accessible peaks were identified using the DiffBind (Stark and Brown, 2011, http://bioconductor.org/packages/release/bioc/vignettes/DiffBind/inst/doc/DiffBind.pdf). For this comparative analysis we used the set of ATAC peaks identified by the ENCODE analysis pipeline using the most restrictive approach (0.05 IDR of true replicates). The default DESEQ2 analysis using a threshold of 0.001 was used to define highly differential ATAC peaks. For each of the three comparisons shown only the peaks identified in the two conditions being compared were considered. For visualization, bigwig files generated by the ENCODE pipeline that represent the p value signal of pooled true replicates were loaded into Integrated Genomics Viewer (IGV). Heatmaps and plot profiles were generated using the plotHeatmap and plotProfile function in DeepTools suite(46). To define the chromatin state of differentially enriched ATAC Peaks the ChromHMM model generated as part of the Roadmap consortium, using H1-derived NPC cells(47), was employed. Briefly, ATAC peaks identified as differentially enriched were intersected with ChromHMM states not taking into account the size of intersect. Bedtools was used for intersections(48) and more than one state could be assigned to the same peak. Intersections with the Quiescent state (characterized by not having enrichment of any chromatin mark) is not shown in the main figure but was assigned to 1066 peaks. Protein binding motifs enriched in differentially accessible enhancer peaks were identified using HOMER(49) by running the −size given and −mask parameters. Motifs with a p-value lower than 1E-12were considered to be significantly enriched. Differentially accessible ATAC peaks were associated with genes using GREAT(50) using the default association tool. Gene ontology for biological processes was also carried out using GREAT. SMARCA4 and H3K27ac chip-seq data (<u>GSE122631</u>) were obtained from(28). ChIP-seq data was aligned to the human genome (hg38) using bowtie2(51) and a MAPQ filter of 10. Duplicate reads were removed using picard tools (http://broadinstitute.github.io/picard/). Enriched regions were called as peaks using MACS2(52) and corresponding input control. The two files of MACS2 peaks for each SMARCA4 ChIP-seq replicates were merged and used in bedtools to calculate total intersection with differentially enriched peaks in shOTUD5 NC3 cells. Intersect size was not taken into account. For visualization in genome browser and heatmap only one randomly selected replicate of SMARCA4 ChIP-seq is shown.

### RNA-seq

RNA from three replicates for cells treated with control shRNA or shRNA against OTUD5, at each stage of neural-conversion, was isolated using Trizol. After confirming that the RNA integration number for each sample was above 8, libraries were prepared using TruSeq Stranded mRNA prep kit with PolyA purificaton and sequenced on HiSeq 2500 using a 1x50 single read mode. RNA-seq analysis and identification of differentially expressed genes was performed using LCDB workflow (https://github.com/lcdb/lcdb-wf). For visualization, bigwig files created by the LCDB workflow were loaded onto IGV.

## Acknowledgments

We would like to thank the patients and their families for participating in research studies. We thank the NHLBI iPS cell core, the NHGRI mouse core, the NICHD molecular genomics core, computational resources of the NIH HPC Biowulf cluster (hpc.nih.gov), and the NIDCR imaging core for excellent technical assistance and Dr. Jacqueline Mays for help with data analysis for teratoma studies. We also would like to thank Dr. Richard Youle for critically reading this manuscript. We would also like to acknowledge the Members of the Undiagnosed Diseases Network (UDN): Maria T. Acosta, Margaret Adam, David R. Adams, Pankaj B. Agrawal, Mercedes E. Alejandro, Patrick Allard, Justin Alvey, Laura Amendola, Ashley Andrews, Euan A. Ashley, Mahshid S. Azamian, Carlos A. Bacino, Guney Bademci, Eva Baker, Ashok Balasubramanyam, Dustin Baldridge, Jim Bale, Michael Bamshad, Deborah Barbouth, Gabriel F. Batzli, Pinar Bayrak-Toydemir, Anita Beck, Alan H. Beggs, Gill Bejerano, Hugo J. Bellen, Jimmy Bennet, Beverly Berg-Rood, Raphael Bernier, Jonathan A. Bernstein, Gerard T. Berry, Anna Bican, Stephanie Bivona, Elizabeth Blue, John Bohnsack, Carsten Bonnenmann, Devon Bonner, Lorenzo Botto, Lauren C. Briere, Elly Brokamp, Elizabeth A. Burke, Lindsay C. Burrage, Manish J. Butte, Peter Byers, John Carey, Olveen Carrasquillo, Ta Chen Peter Chang, Sirisak Chanprasert, Hsiao-Tuan Chao, Gary D. Clark, Terra R. Coakley, Laurel A. Cobban, Joy D. Cogan, F. Sessions Cole, Heather A. Colley, Cynthia M. Cooper, Heidi Cope, William J. Craigen, Michael Cunningham, Precilla D’Souza, Hongzheng Dai, Surendra Dasari, Mariska Davids, Jyoti G. Dayal, Esteban C. Dell’Angelica, Shweta U. Dhar, Katrina Dipple, Daniel Doherty, Naghmeh Dorrani, Emilie D. Douine, David D. Draper, Laura Duncan, Dawn Earl, David J. Eckstein, Lisa T. Emrick, Christine M. Eng, Cecilia Esteves, Tyra Estwick, Liliana Fernandez, Carlos Ferreira, Elizabeth L. Fieg, Paul G. Fisher, Brent L. Fogel, Irman Forghani, Laure Fresard, William A. Gahl, Ian Glass, Rena A. Godfrey, Katie GoldenGrant, Alica M. Goldman, David B. Goldstein, Alana Grajewski, Catherine A. Groden, Andrea L. Gropman, Sihoun Hahn, Rizwan Hamid, Neil A. Hanchard, Nichole Hayes, Frances High, Anne Hing, Fuki M. Hisama, Ingrid A. Holm, Jason Hom, Martha Horike-Pyne, Alden Huang, Yong Huang, Rosario Isasi, Fariha Jamal, Gail P. Jarvik, Jeffrey Jarvik, Suman Jayadev, Yong-hui Jiang, Jean M. Johnston, Lefkothea Karaviti, Emily G. Kelley, Dana Kiley, Isaac S. Kohane, Jennefer N. Kohler, Deborah Krakow, Donna M. Krasnewich, Susan Korrick, Mary Koziura, Joel B. Krier, Seema R. Lalani, Byron Lam, Christina Lam, Brendan C. Lanpher, Ian R. Lanza, C. Christopher Lau, Kimberly LeBlanc, Brendan H. Lee, Hane Lee, Roy Levitt, Richard A. Lewis, Sharyn A. Lincoln, Pengfei Liu, Xue Zhong Liu, Nicola Longo, Sandra K. Loo, Joseph Loscalzo, Richard L. Maas, Ellen F. Macnamara, Calum A. MacRae, Valerie V. Maduro, Marta M. Majcherska, May Christine V. Malicdan, Laura A. Mamounas, Teri A. Manolio, Rong Mao, Kenneth Maravilla, Thomas C. Markello, Ronit Marom, Gabor Marth, Beth A. Martin, Martin G. Martin, Julian A. Martínez-Agosto, Shruti Marwaha, Jacob McCauley, Allyn McConkie-Rosell, Colleen E. McCormack, Alexa T. McCray, Heather Mefford, J. Lawrence Merritt, Matthew Might, Ghayda Mirzaa, Eva Morava-Kozicz, Paolo M. Moretti, Marie Morimoto, John J. Mulvihill, David R. Murdock, Avi Nath, Stan F. Nelson, John H. Newman, Sarah K. Nicholas, Deborah Nickerson, Donna Novacic, Devin Oglesbee, James P. Orengo, Laura Pace, Stephen Pak, J. Carl Pallais, Christina GS. Palmer, Jeanette C. Papp, Neil H. Parker, John A. Phillips III, Jennifer E. Posey, John H. Postlethwait, Lorraine Potocki, Barbara N. Pusey, Aaron Quinlan, Wendy Raskind, Archana N. Raja, Genecee Renteria, Chloe M. Reuter, Lynette Rives, Amy K. Robertson, Lance H. Rodan, Jill A. Rosenfeld, Robb K. Rowley, Maura Ruzhnikov, Ralph Sacco, Jacinda B. Sampson, Susan L. Samson, Mario Saporta, C. Ron Scott, Judy Schaechter, Timothy Schedl, Kelly Schoch, Daryl A. Scott, Lisa Shakachite, Prashant Sharma, Vandana Shashi, Jimann Shin, Rebecca Signer, Catherine H. Sillari, Edwin K. Silverman, Janet S. Sinsheimer, Kathy Sisco, Kevin S. Smith, Lilianna Solnica-Krezel, Rebecca C. Spillmann, Joan M. Stoler, Nicholas Stong, Jennifer A. Sullivan, Angela Sun, Shirley Sutton, David A. Sweetser, Virginia Sybert, Holly K. Tabor, Cecelia P. Tamburro, Queenie K.-G. Tan, Mustafa Tekin, Fred Telischi, Willa Thorson, Cynthia J. Tifft, Camilo Toro, Alyssa A. Tran, Tiina K. Urv, Matt Velinder, Dave Viskochil, Tiphanie P. Vogel, Colleen E. Wahl, Stephanie Wallace, Nicole M. Walley, Chris A. Walsh, Melissa Walker, Jennifer Wambach, Jijun Wan, Lee-kai Wang, Michael F. Wangler, Patricia A. Ward, Daniel Wegner, Mark Wener, Monte Westerfield, Matthew T. Wheeler, Anastasia L. Wise, Lynne A. Wolfe, Jeremy D. Woods, Shinya Yamamoto, John Yang, Amanda J. Yoon, Guoyun Yu, Diane B. Zastrow, Chunli Zhao, Stephan Zuchner.

## Author contributions

D.B.B. conceived and designed the study, identified and saw OTUD5 patients, wrote the manuscript and together with M.A.B. performed most experiments. M.A.B. designed, performed, and interpreted stem cell differentiation, in vitro deubiquitylation, and immunoprecipitation experiments. A.J.A anlaysed and interpreted all mass spectrometry experiments and performed all immunofluorescence exerpiments. J.T. performed and analyzed ATAC seq and RNA seq experiments and wrote the manuscript. H.O. analyzed RNA seq experiments. R.D. and A.M. analyzed ATAC seq and RNA seq experiments. D.T.U. performed and analyzed exome sequencing in and methylation studies in P6-F4, and J.I. supervised the project. K.S., E.M., P.D., J.B, W. M., K.B., N.M., R.W. M.K., Y.N., S.O., T.K., N.M., M.W., and C.J.T. helped provide clinical information on patients reported in this study. I.A. conceived and designed the study and interpreted results. D.L.K. conceived and designed the study, interpreted results and secured funding. P.P.R. designed, performed, and analyzed ATAC seq and RNA seq experiments, wrote the manuscript, and secured funding. A.W. conceived and designed the study, performed experiments, interpreted results, wrote the manuscript, and secured funding.

## Supplementary Materials

Figures S1-S9

Tables S1-S5

**Fig. S1:**
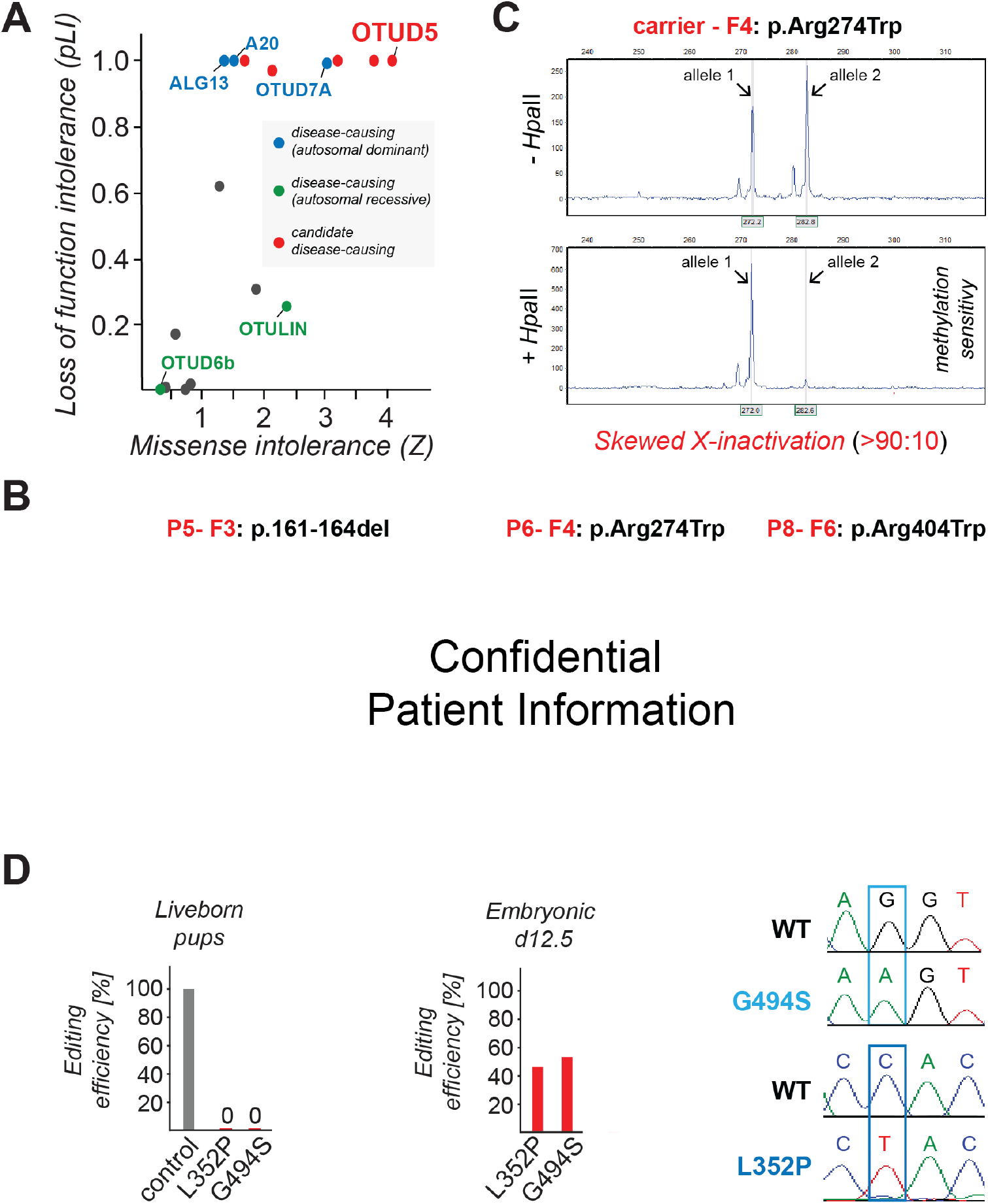
OTUD5 is essential for proper human and mouse development. (**A**) Amongst several OTU DUBs, OTUD5 is the strongest candidate for hypomorphic mutations leading to disease given high loss of function and missense intolerance scores. Loss of function intolerance (pLI) and missense intolerance (Z) were determined for all OTU DUBs using gnomAD. (**B**) Clinical photos showing craniofacial (retrognathia, midface hypoplasia, hypertelorism, low set posteriorly rotated ears) of patient P5-F3 carrying the p.161-164del mutation, patient P6-F4 carrying the p.Arg274Trp mutation, or patient P8-F6 carrying the p.Arg404Trp mutation. (**C**) The *OTUD5* p.Arg274Trp carrier mother exhibits skewed X-inactivation, as revealed by digestion of genomic DNA with a the methylation-sensitive restriction enzyme *HpaII* followed by PCR amplification of the human androgen receptor (*AR*) gene at Xq12 and capillary gel electrophoresis. DNA was isolated from peripheral blood. (**D**) CRISPR-mediated knock-out of *OTUD5* or knock-in of the p.Gly494Ser or p.Leu352Pro patient variants results in embryonic lethality. *Left graph:* Mouse zygotes were injected with Cas9 complexed with guide RNAs and respective repair oligos and transferred into pseudo pregnant recipient mice. Percentage of liveborn pups with edited alleles (knock out or knock in) for a non-essential gene (control), OTUD5^L352P^, or OTUD5^G494S^ is shown (n>70 injected embryos per condition) *Right graph*: Mouse embryos were injected with guide RNA loaded Cas9 and respective repair oligos and implanted into mice. Pregnant mice were sacrificed and embryos isolated at day E12.5. Percentage of pups with edited alleles (knock out or knock in) for *OTUD5^L352P^* or *OTUD5^G494S^* are shown (n>70 injected embryos per condition). Sanger sequencing depicting examples of E12.5 knock-in embryos are shown.

**Fig. S2:**
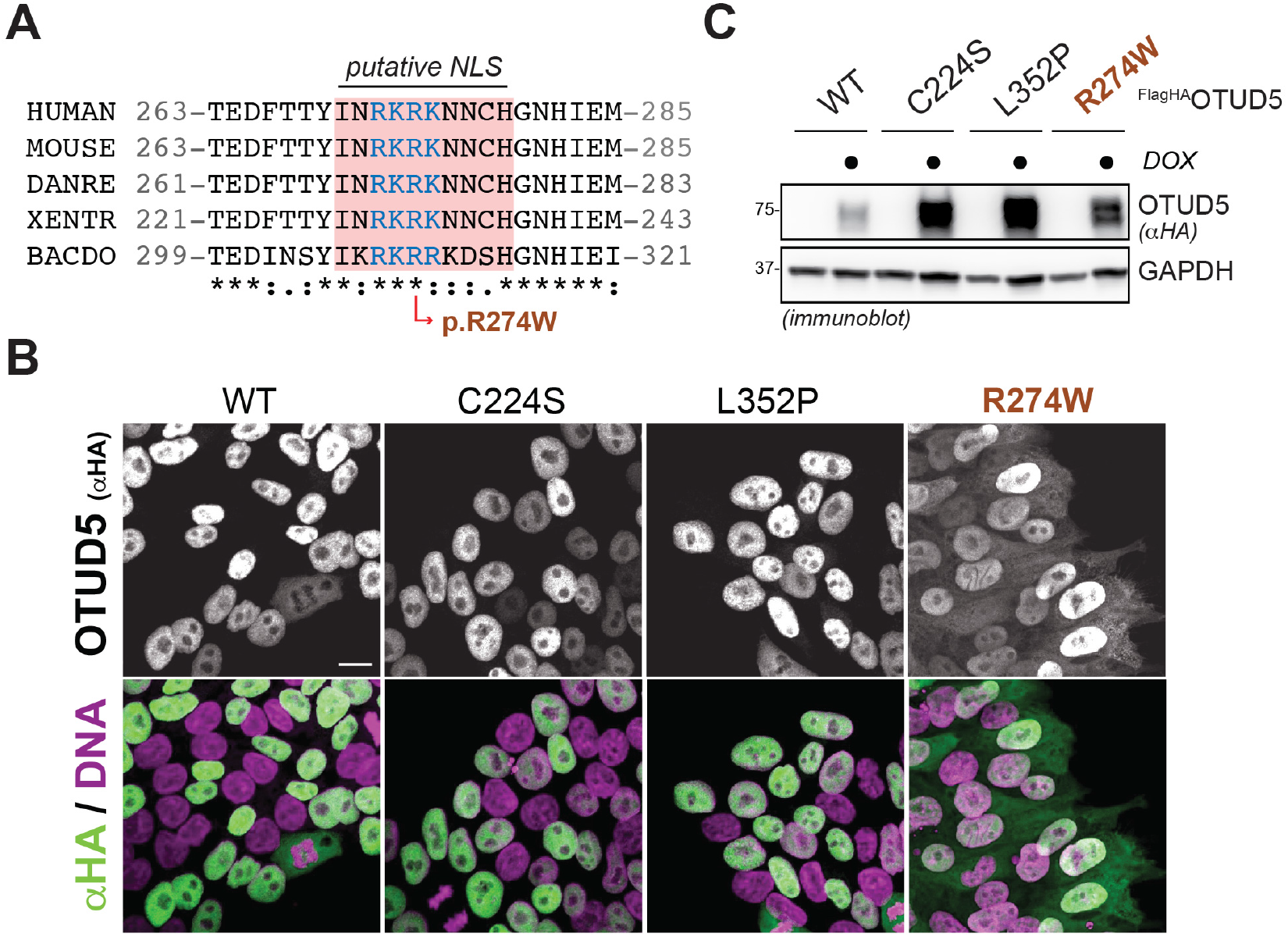
The p.Arg274Trp OTUD5 patient mutation is present in a putative NLS and results in protein mislocalization. (**A**) The p.Arg274Trp (R274W) patient variant is present in a putative NLS that is conserved amongst species. Sequence alignment of OTUD5 was performed using clustal omega. (**B**) The OTUD5 R274W mutant protein partially mislocalizes to the cytoplasm. hES H1 cells stably expressing doxycycline-inducible wildtype ^FLAGHA^OTUD5 (WT), catalytically inactive ^FLAGHA^OTUD5 (C224S), or patient variant ^FLAGHA^OTUD5 (L352P, R274W) were induced with doxycycline (DOX) for 48h, fixed, and subjected to anti-HA immunofluorescence microscopy (C) ^FLAGHA^OTUD5^R274W^ does not express higher than wildtype ^FLAGHA^OTUD5. hES H1 cells stably expressing doxycycline-inducible wildtype ^FLAGHA^OTUD5 (WT), catalytically inactive ^FLAGHA^OTUD5 (C224S), or patient variant ^FLAGHA^OTUD5 (L352P, R274W) were induced with doxycycline (DOX) for 48h as indicated and subjected to anti-HA and anti-GAPDH immunoblot analysis.

**Fig. S3:**
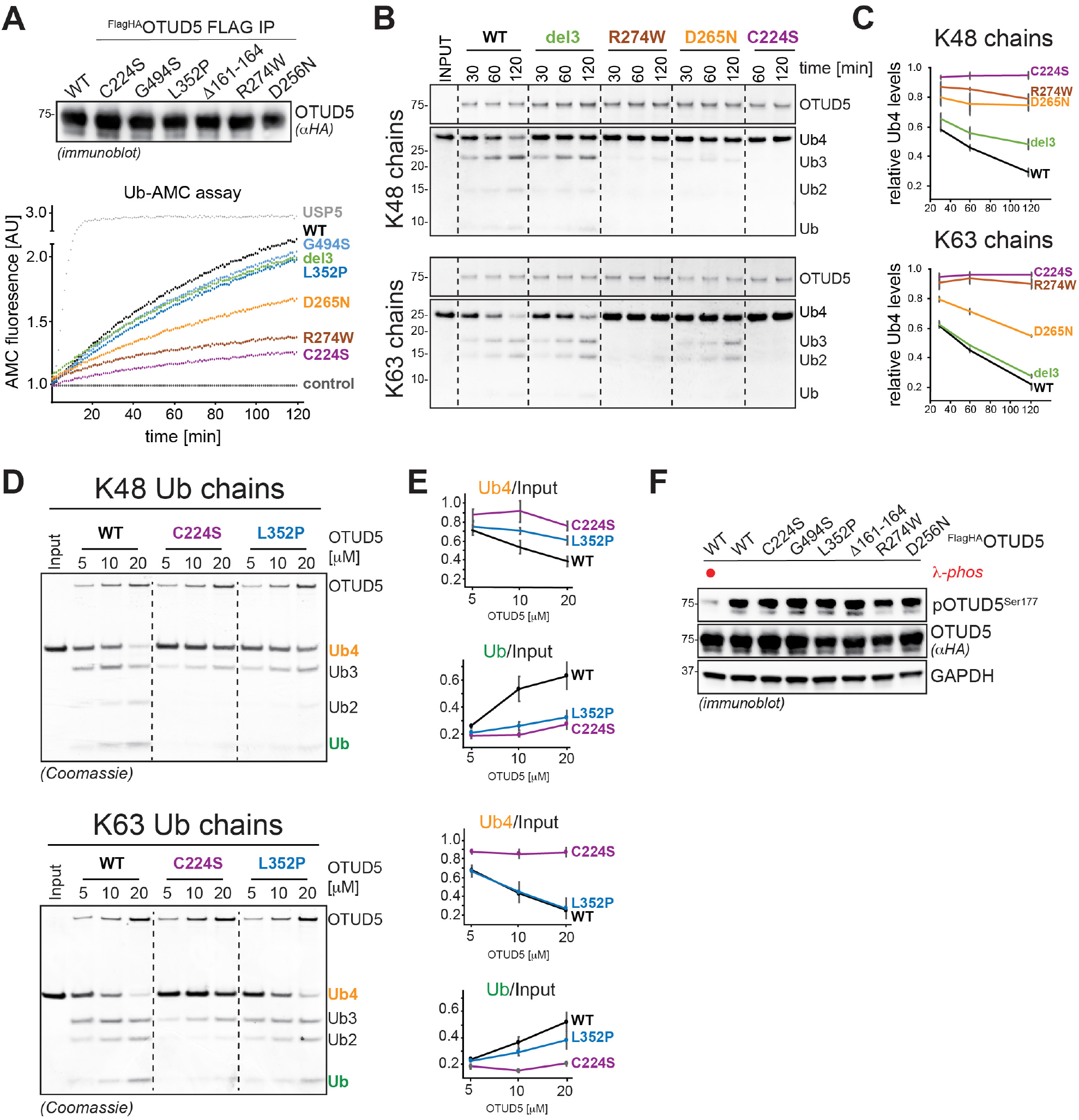
Most patient mutations reduce OTUD5 deubiquitylation activity. (**A**) The p.D265N and p.R274W mutations reduce OTUD5’s ability to hydrolyze the model DUB substrate Ub-AMC, while other variants have no significant effect. *Upper panel*: Wildtype ^FLAGHA^OTUD5 (WT), catalytically inactive ^FLAGHA^OTUD5 (C224S), and denoted patient variant ^FLAGHA^OTUD5 were purified from HEK293T cells followed by normalization and anti-HA immunoblot analysis. *Lower panel*: Purified ^FLAGHA^OTUD5 variants and ^FLAGHA^USP5 were incubated with Ub-AMC and increase of AMC fluorescence was detected over time. Control = flag-IPs from HEK293T cells. (**B**) The p.D265N and p.R274W variants reduce OTUD5 cleavage activity towards K63- and K48-chains, while the p.161-164del variant specifically reduces K48-ubiquitin chain cleavage. Wildtype ^FLAGHA^OTUD5 (WT), catalytically inactive ^FLAGHA^OTUD5 (C224S), and patient variant ^FLAGHA^OTUD5 (Del, R274W, or D265N) were purified from HEK293T cells and incubated with tetra-K48- or tetra-K63-ubiquitin chains for indicated time periods and analyzed by colloidal coomassie-stained SDS PAGE gels. (**C**) Quantification of three independent *in vitro* deubiquitylation experiments shown in panel b (error bars denote s.e.m). Intensity of Ub4 band is relative to the sum of intensity of Ub3, Ub2, and Ub band. (**D**) The p.L352P variant specifically reduces OTUD5’s K48-ubiquitin chain cleavage activity. Wildtype ^FLAGHA^OTUD5 (WT), catalytically inactive ^FLAGHA^OTUD5 (C224S), and patient variant ^FLAGHA^OTUD5 (L352P) were purified from HEK293T cells, quantified, and increasing amounts of enzyme were incubated with K48-linked or K63-linked tetra-ubiquitin chains (Ub4) for 2h. Samples were analyzed by colloidal coomassie-stained SDS PAGE gels. (**E**) Quantification of three independent *in vitro* deubiquitylation experiments shown in panel d (error bars denote s.e.m). Intensities of Ub4 or Ub bands are relative to intensity of Ub4 input band (**F**) Patient variants have no significant impact on OTUD5’s activating phosphorylation. Wildtype ^FLAGHA^OTUD5 (WT), catalytically inactive ^FLAGHA^OTUD5 (C224S), and denoted patient variant ^FLAGHA^OTUD5 were expressed in HEK293T cells. Cells were lysed and treated with λ-phosphatase as indicated followed by anti-HA, anti-pOTUD5^Ser177^, and anti-GAPDH immunoblot analysis.

**Fig. S4:**
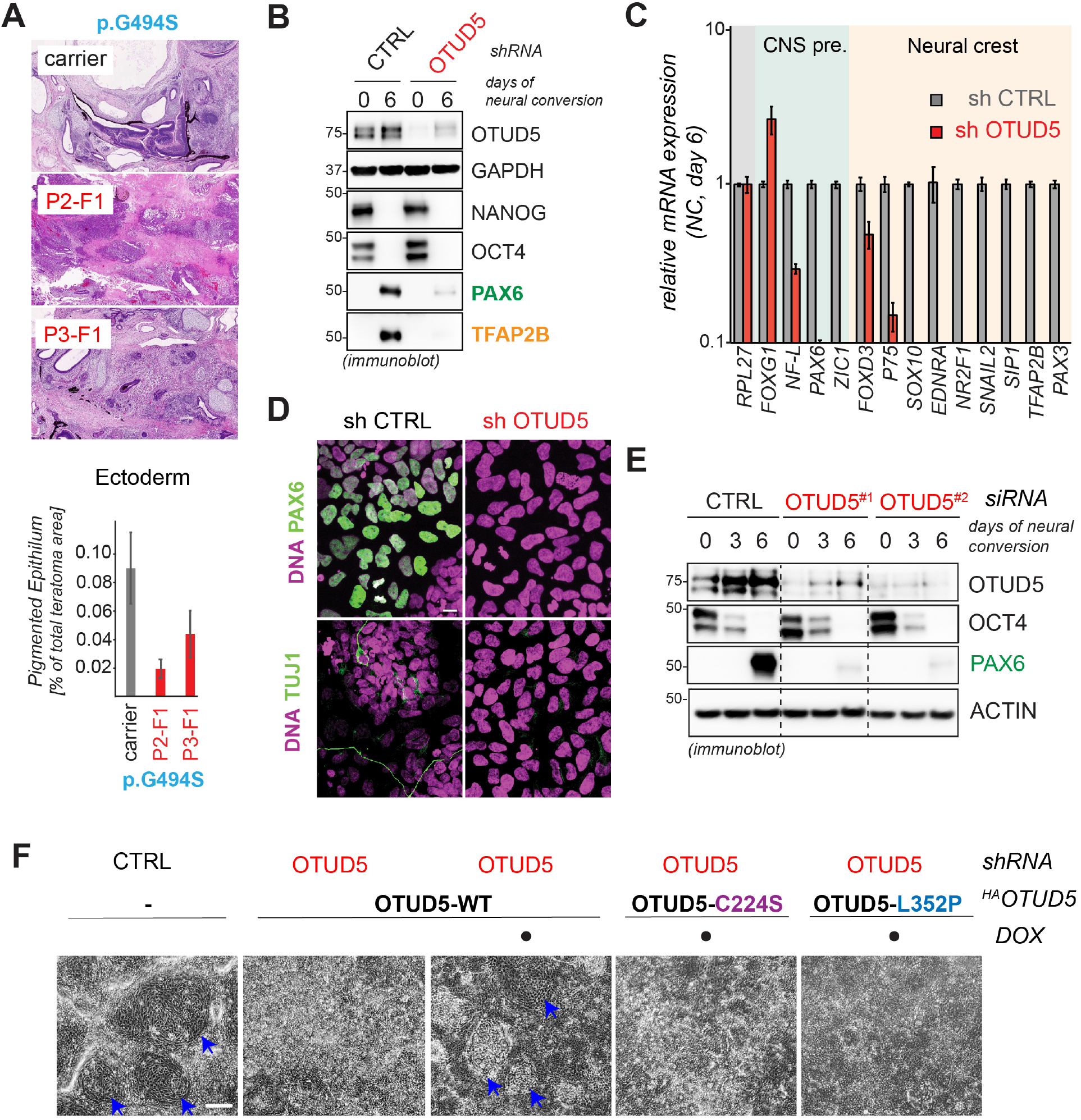
Reduction of OTUD5 activity results in aberrant neuro-ectodermal differentiation. (**A**) iPSCs derived from OTUD5 p.G494S patients are impaired in neuroectodermal differentiation *in vivo*. iPSCs derived from two sibling patients with p.Gly494Ser or their mother carrier were injected into immunocompromised mice and allowed to develop into teratomas for 8 weeks. Teratomas were isolated, sectioned, and stained with hematoxylin and eosin. Graph depicts quantification of the area occupied by pigmented epithelium (ectodermal marker) and quantified using image J (error bars denote s.e.m, 12 slides of 2 teratomas were used per condition. * = p <0.05, unpaired t-test). (**B**) Depletion of OTUD5 from hES H1 cells causes aberrant neural conversion. hES H1 cells were depleted of endogenous OTUD5 using stably expressed shRNA and subjected to neural conversion for 6 days. Success of cell differentiation was monitored by immunoblotting using antibodies against NANOG and OCT4 (hESC markers), PAX6 (CNS precursor marker), and TFAP2B (neural crest marker) and GAPDH (loading control). (**C**) Depletion of OTUD5 from hES H1 cells causes aberrant neural conversion. Control or OTUD5-depleted hES H1 cells were subjected to neural conversion for 6 days and analyzed by qRT-PCR for expression of CNS precursor markers (highlighted in green) and neural crest markers (highlighted in orange). Marker expression was normalized to carrier control followed by hierarchical cluster analysis. RPL27 was used as endogenous control (n=3 technical replicates, error bars denote s.e.m.) (**D**) Depletion of OTUD5 from hES H1 cells causes aberrant neural conversion, as seen at the single cell level. Control or OTUD5-depleted hES H1 cells were subjected to neural conversion for 9d and the success of differentiation was determined by immunofluorescence microscopy using antibodies against PAX6 (CNS precursor marker), TUJ1 (neuronal marker). Scale Bar = 20μm. (**E**) Depletion of OTUD5 from hES H1 cells causes aberrant neural conversion as show at the protein level. hES H1 cells were depleted of endogenous OTUD5 using two different pools of siRNA and subjected to neural conversion for 3 and 6 days. Success of cell differentiation was monitored by immunoblotting using antibodies against OCT4 (hESC marker) and PAX6 (CNS precursor marker) and ACTIN (loading control). (**F**) K48-chain specific deubiquitylation activity of OTUD5 is required for proper CNS precursor and neural crest differentiation. hES H1 cells stably expressing shRNA-resistant and doxycycline-inducible wildtype (WT), catalytically inactive (C224S), or K48-chain cleavage deficient (L352P) ^HA^OTUD5 were generated. Cells were then depleted of endogenous OTUD5 using shRNA as indicated, treated with or without doxycycline (DOX), and subjected to neural conversion for 6 days. Success of differentiation was determined by detection of neural rosette structures using phase contrast microscopy. Neural rosettes are highlighted by blue arrows. This experiment was also analyzed by immunoblotting as depicted in Fig. 2E.

**Fig. S5:**
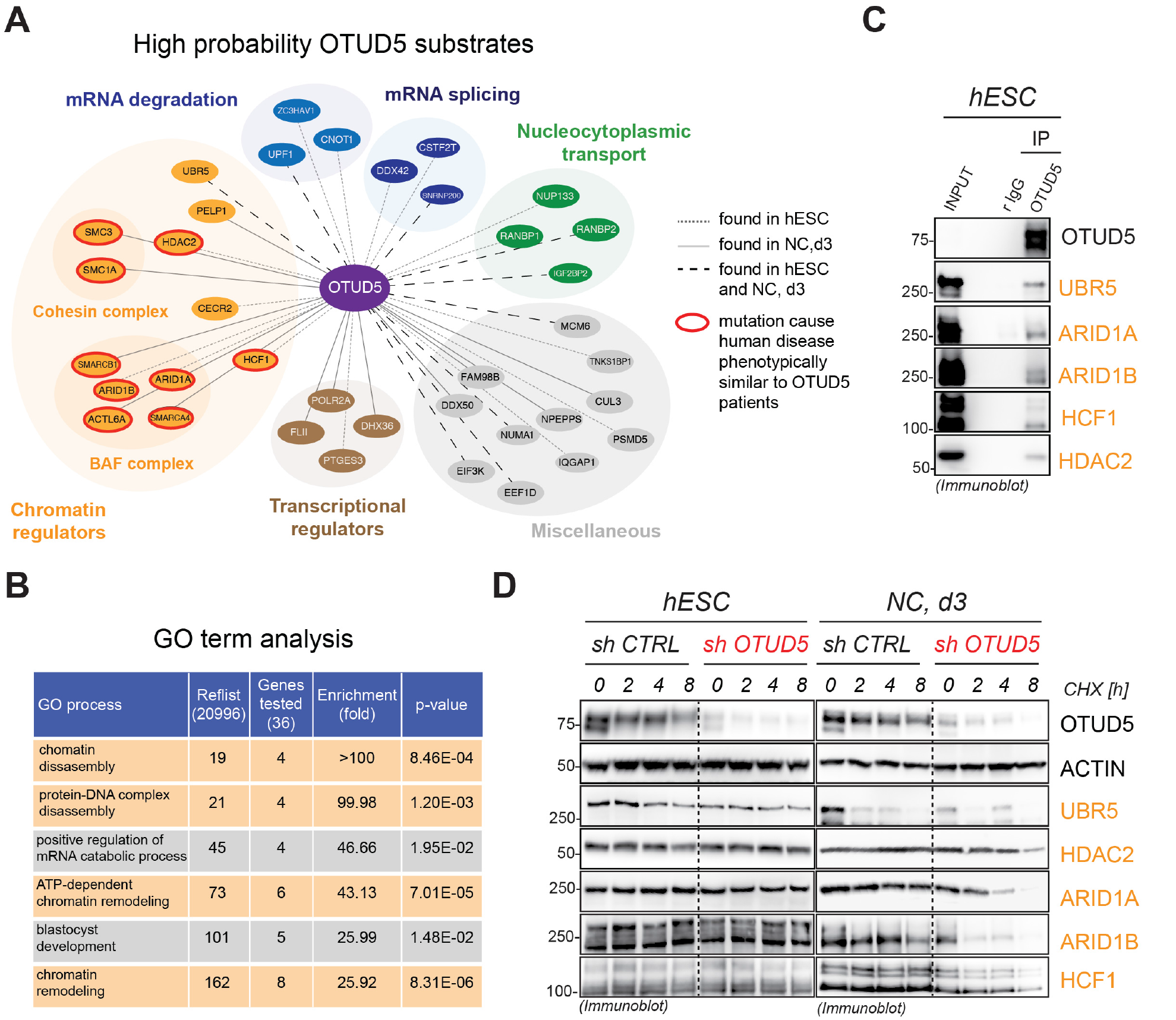
OTUD5 interacts with chromatin regulators and regulates their stability specifically during differentiation. (**A**) Schematic representation of high probability substrates of OTUD5. To identify high probability substrates of OTUD5 two independent proteomic experiments were performed (cf. Figure 3a). First, control or OTUD5-depleted H1 hESCs or hESCs undergoing neural conversion for 1 or 3 days were lysed and ubiquitylated proteins were isolated by TUBE pull down followed by protein identification via mass spectrometry. Second, self-renewing or differentiating control hESCs or hESCs expressing wildtype (WT) or catalytically inactive (C224S) ^FLAG^OTUD5 were lysed and subjected to anti-FLAG immunoprecipitation followed by identification of interacting proteins via mass spectrometry. Candidate OTUD5 substrates were defined as proteins found to be more ubiquitylated upon OTUD5 depletion and identified as specific OTUD5 WT or C224S interactors. These high probability substrates of OTUD5 were functionally annotated and plotted using cytoscape. Color of lines indicate in which cell state the physical OTUD5 interaction occurred (dark grey: hES cell state (hESC), light grey: cells undergoing neural conversion for 3d (NC, d3), black: hESC and NC, d3). Candidate substrates that when mutated cause human diseases with phenotypic overlap to OTUD5 patients are circled in red. (**B**) High probability substrates of OTUD5 are significantly enriched in chromatin regulators, as determined by GO term analysis. (**C**) OTUD5 endogenously interacts with chromatin regulators. hES H1 cells (5 x 15cm plates per condition) were lysed and lysates were subjected to anti-OTUD5 immunoprecipitiation followed by SDS PAGE and immunoblot analysis using indicated antibodies. Rabbit IgGs were used as control. (**D**) OTUD5 stabilizes chromatin regulators in differentiating, but not self-renewing hESCs. Control or OTUD5-depleted hESCs or hESC subjected to neural conversion for 3 days were treated with cycloheximide for indicated time periods and protein stability of HDAC2, UBR5, ARID1A/B, and HCFC1 was determined by immunoblotting using indicated antibodies. A representative experiment of a total of three biological replicates is shown. Quantification is depicted in Fig. 3D.

**Fig. S6:**
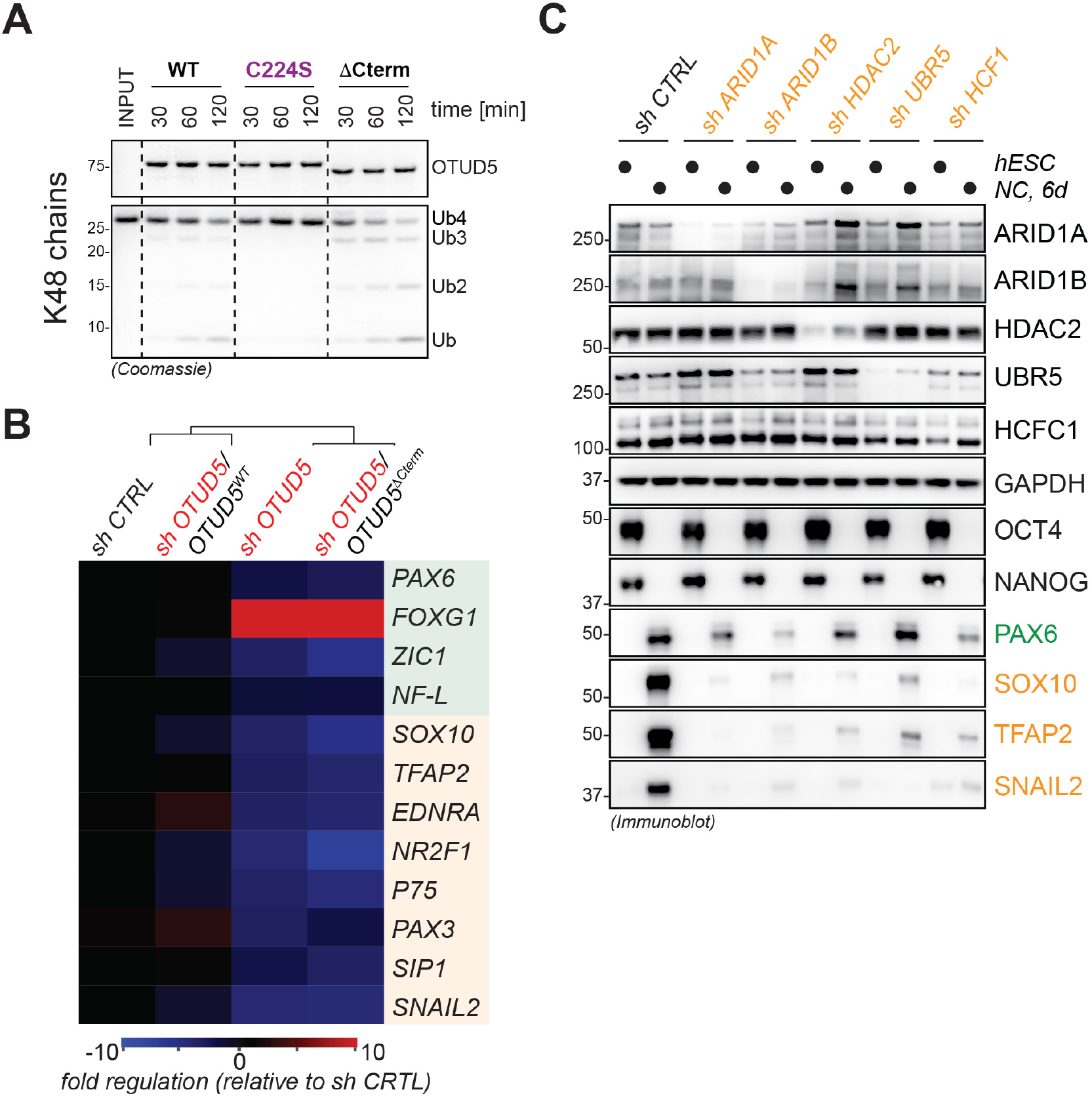
OTUD5 regulates differentiation through stabilizing chromatin remodelers. (**A**) Deletion of the C-terminus of OTUD5 containing the UIM motif does not reduce its K48-ubiquitin chain activity *in vitro*. Wildtype ^FLAGHA^OTUD5 (WT), catalytically inactive ^FLAGHA^OTUD5 (C224S), and chromatin regulator binding-deficient OTUD5 (ΔCterm) were purified from HEK293T cells and incubated with tetra-K48-chains for indicated time periods and analyzed by colloidal coomassie-stained SDS PAGE gels. (B) The chromatin regulator binding-deficient ^FLAGHA^OTUD5^ΔCterm^ mutant does not support neural conversion. hES H1 cells stably expressing shRNA-resistant and doxycycline-inducible wildtype (WT) or chromatin regulator binding-deficient (ΔCterm) ^FLAGHA^OTUD5. Cells were then depleted of endogenous OTUD5 using shRNA as indicated, treated with or without doxycycline (DOX), and subjected to neural conversion for 6 days. This was followed by qRT-PCR for expression of CNS precursor markers (highlighted in green) and neural crest markers (highlighted in orange). Marker expression was normalized to carrier control followed by hierarchical cluster analysis. RPL27 was used as endogenous control. (C) Depletion of chromatin regulators results in aberrant neural conversion. hES H1 cells were depleted of endogenous indicated chromatin regulators using stably expressed shRNAs and subjected to neural conversion for 6 days and analyzed by immunoblotting to assess knockdown efficiency and the expression of CNS precursor marker (PAX6) and neural crest markers (SOX10, TFAP2, SNAIL2). GAPDH serves as loading control.

**Fig. S7:**
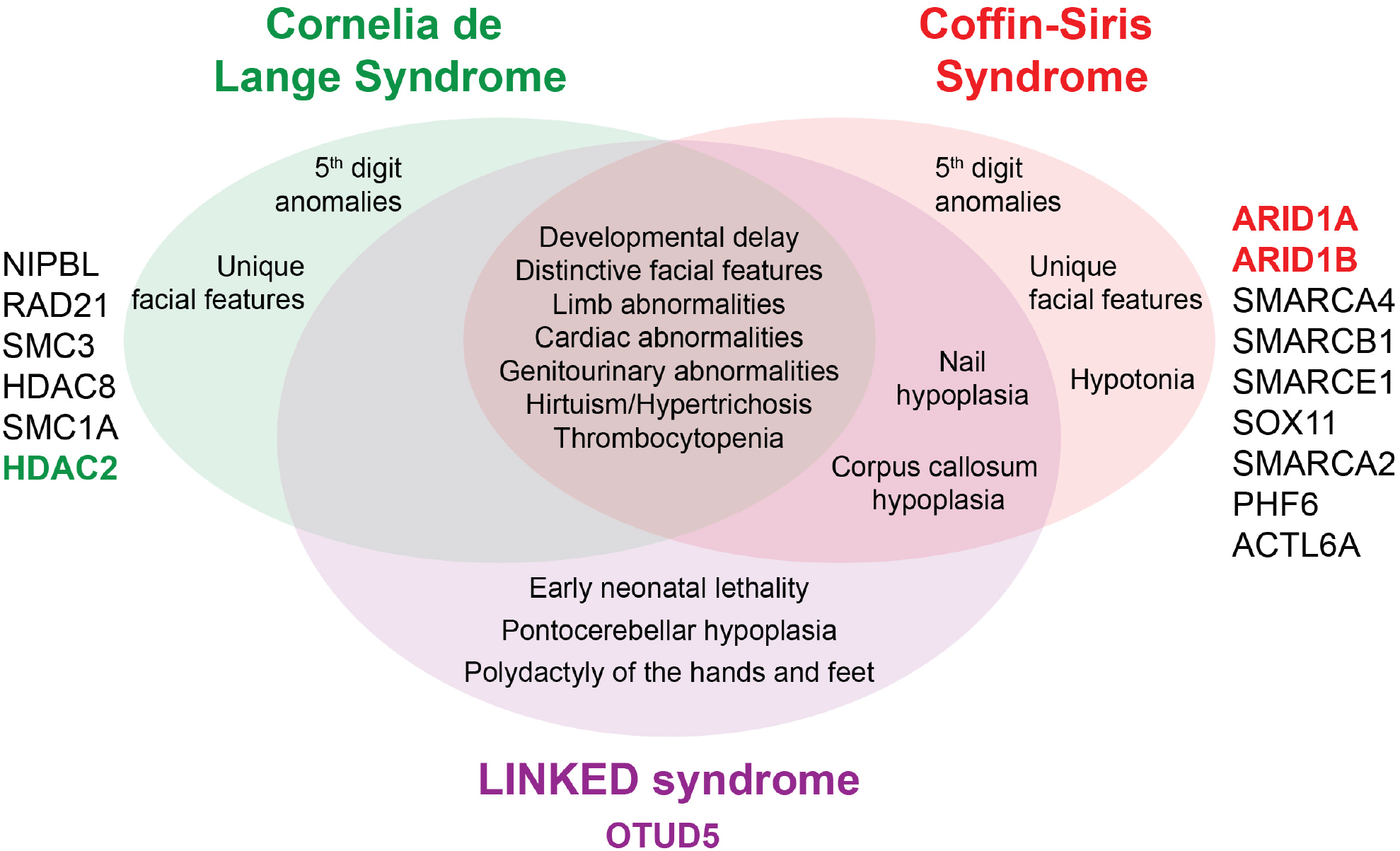
Venn Diagram depicting the unique and overlapping features of LINKED patients and patients suffering from Cornelia de Lange and Coffin-Siris Syndrome. Genes whose mutations cause the respective diseases are depicted. HDAC2 and ARID1A/B, substrates of OTUD5, are highlighted in bold.

**Fig. S8:**
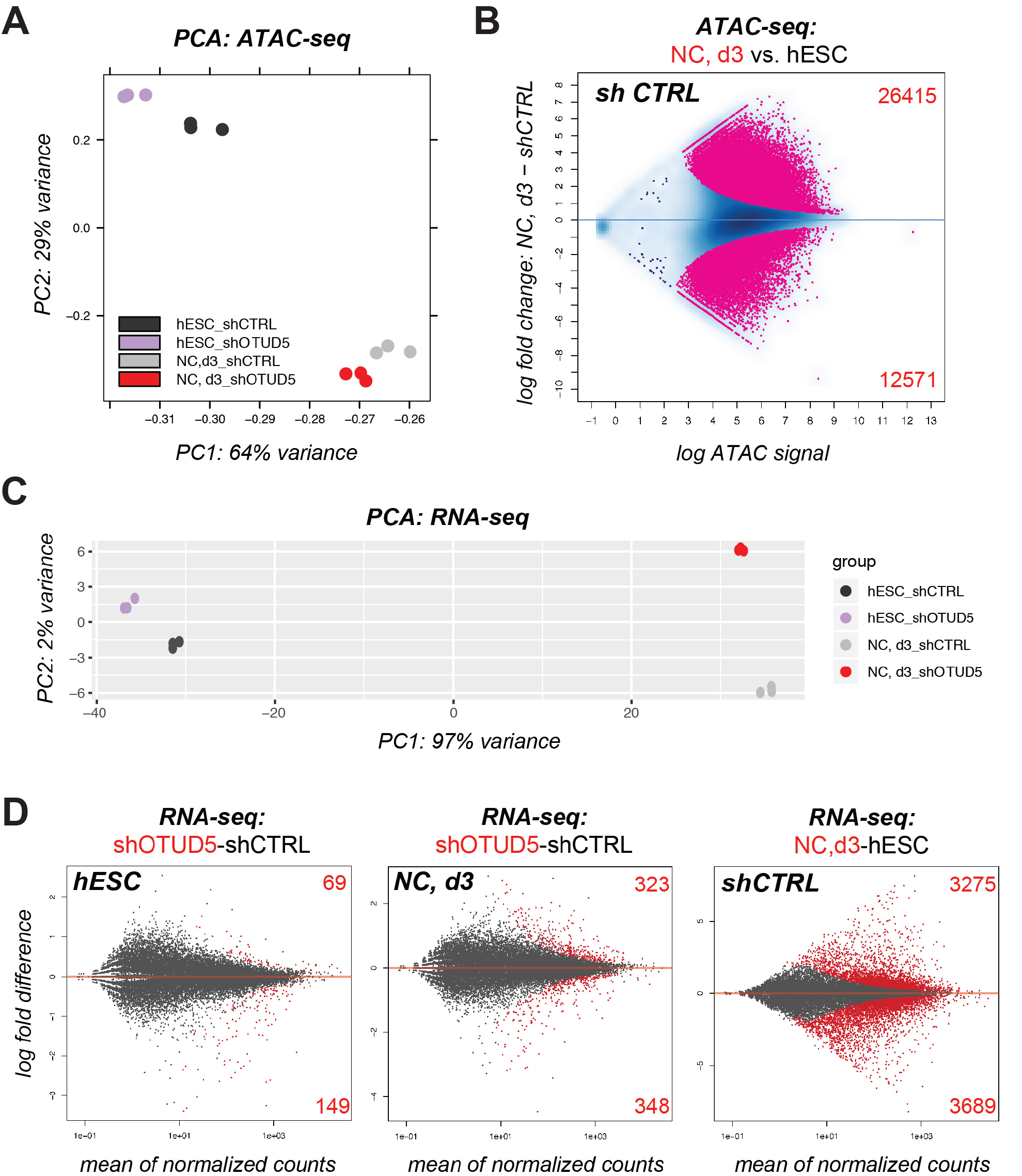
Chromatin accessibility and transcriptome remain largely unchanged upon loss of OTUD5 during early differentiation. (**A**) Principal component analysis (PCA) of chromatin accessibility (assayed by ATAC-seq) of control and OTUD5-depleted hES H1 cells (hESC) and cells subjected to neural conversion for three days (NC, d3). OTUD5 has only minor effects on the overall chromatin landscape as compared to the differences observed during neural conversion (hESC versus NC3,d). (**B**) MAplot highlighting the drastically different ATAC-seq profiles between self-renewing hES H1 cells (hESC) and cells undergoing neural conversion for 3 days (NC, d3). Peaks represented in the top part of the plot gain accessibility during differentiation while peaks in the bottom half, lose accessibility. Pink dots represent peaks with statistically significant enrichment differences (adj pvalue < 0.0001) between the two conditions. Note that these changes are drastically more different than differences observed comparing control and OTUD5-depleted cells at NC, d3 in Figure 4a, suggesting that OTUD5-dependent chromatin accessibility changes are not a mere consequence of failed differentiation. (**C**) PCA of transcriptional profiles, assayed using RNAseq, in control and OTUD5-depleted hESC and cells subjected to neural conversion for 3 days (NC, d3). OTUD5 depletion has little effect on transcriptional state of hESCs as compared to the difference in profiles evident at NC, d3 stage. 97% of the transcriptional variance is related to differentiation. (**D**) MA-plots illustrating the minimal impact of OTUD5 loss on transcription in hESCs (left panel), which is slightly increased at day 3 of neural conversion (NC, d3, middle panel). OTUD5-dependent transcriptional changes at hESC (left panel), and NC, d3 (center panel) are modest compared to transcriptional alterations during differentiation in control cells (right panel).

**Fig. S9:**
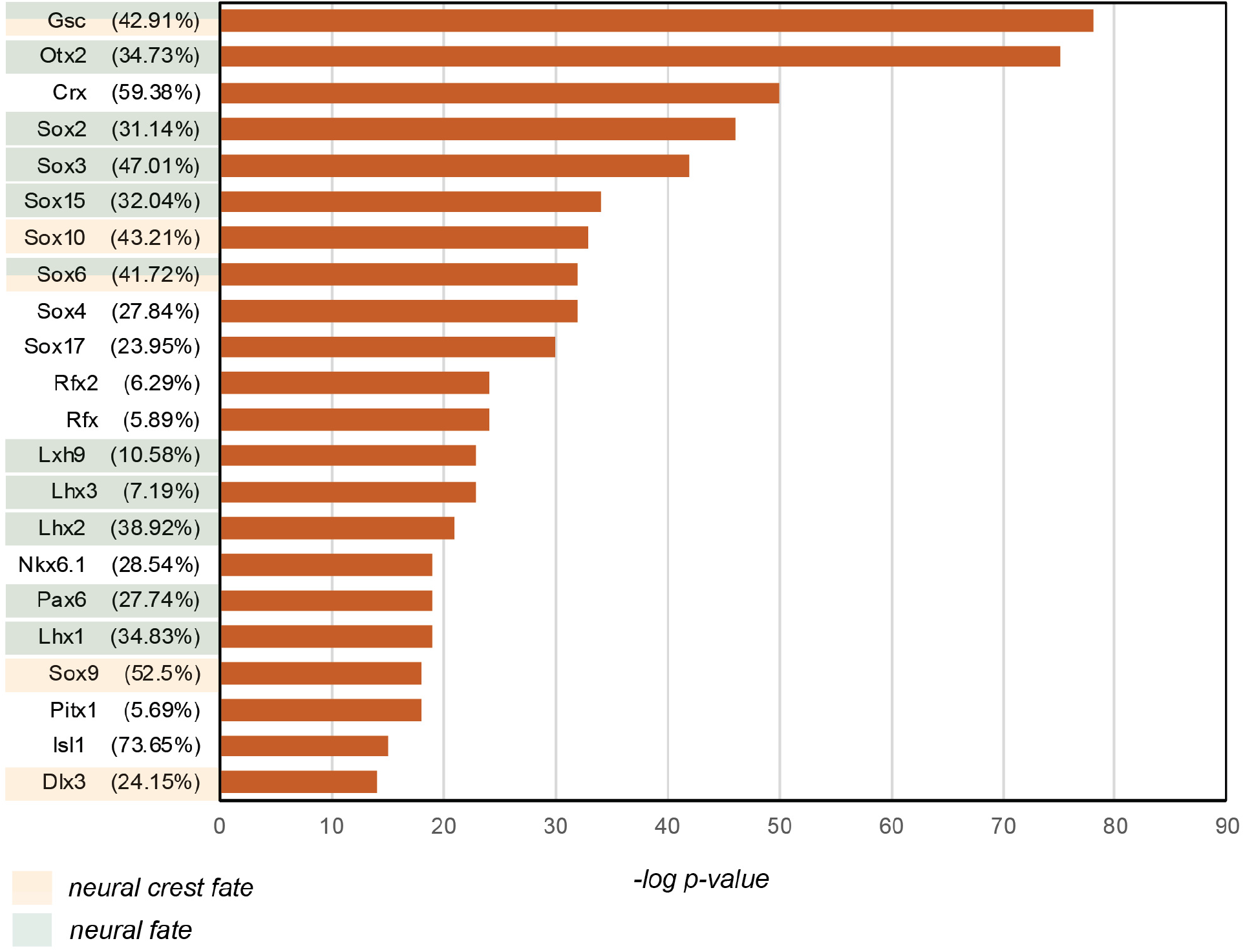
OTUD5-regulated enhancers are enriched in motifs for neural- and neural-crest promoting transcription factors. Transcription factor motif analysis was performed on the ATAC-seq regions that were labeled as potential enhancers and that lost chromatin accessibility upon OTUD5 loss at day 3 of neural conversion. These regions are enriched for transcription factors important for driving differentiation towards a neural fate (highlighted in green) or neural crest fate (highlighted in orange). Numbers in parenthesis represent the fraction of enhancers that contain the respective transcription factor motifs.

**Table S1: Detailed clinical information on OTUD5-deficient patients**

**Table S2: Proteins differentially binding to TUBEs upon OTUD5 depletion**:

Nuclear proteins found more than 5-fold regulated upon OTUD5 depletion in TUBE pull downs from hESCs or hESC undergoing neural conversion for 1d (NC1) or 3 days (NC3) with at least 13 unique peptides and identified in at least 4 conditions (groups) are shown.

**Table S3: Proteins physically interacting with OTUD5 in self renewing and differentiating hESCs.**

Relative Total spectral counts (rTSCs) of nuclear proteins identified in flag-IPs from hESCs or hESC undergoing neural conversion for 3 days (NC3) expressing FLAG-OTUD5 WT or C224S. Only nuclear proteins that were not detected in control IPs (FLAG IPs from hESCs or hESCs undergoing neural conversions for 3 days) are shown.

**Table S4: High probability targets of OTUD5**.

Nuclear proteins found more than 5-fold upregulated upon OTUD5 depletion in TUBE pull downs and specifically interacting with OTUD5 WT or C224S.

**Table S5: qRT-PCR primers used in this study**

